# Emergence of synchronized multicellular mechanosensing from spatiotemporal integration of heterogeneous single-cell information transfer

**DOI:** 10.1101/2020.09.28.316240

**Authors:** Amos Zamir, Guanyu Li, Katelyn Chase, Robert Moskovitch, Bo Sun, Assaf Zaritsky

## Abstract

We quantitatively characterize how noisy and heterogeneous behaviors of individual cells are integrated across a population toward multicellular synchronization by studying the calcium dynamics in mechanically stimulated monolayers of endothelial cells. We used information-theory to quantify the asymmetric information-transfer between pairs of cells and define quantitative measures of how single cells receive or transmit information in the multicellular network. We find that cells take different roles in intercellular information-transfer and that this heterogeneity is associated with synchronization. Cells tended to maintain their roles between consecutive cycles of mechanical stimuli and reinforced them over time, suggesting the existence of a cellular “memory” in intercellular information transfer. Interestingly, we identified a subpopulation of cells characterized by higher probability of both receiving and transmitting information. These “communication hub” roles were stable - once a cell switched to a “communication hub” role it was less probable to switch to other roles. This stableness property of the cells led to gradual enrichment of communication hubs that was associated with the establishment of synchronization. Our analysis demonstrated that multicellular synchronization was established by effective information spread from the (local) single cell to the (global) group scale in the multicellular network. Altogether, we suggest that multicellular synchronization is driven by single cell communication properties, including heterogeneity, functional memory and information flow.

## Introduction

Synchronized multicellular dynamics is the basis of many critical physiological processes, such as the rhythmic beating of cardiomyocytes, planar cell polarity, regulation of vascular tone, and brain activities. However, a fundamental question remains elusive: how synchronization in the group emerges from the interactions of individual cells, each making stochastic decisions based on noisy cues from their local environment?

Every cell influence and respond to cells in its community through a complex interplay of chemical and physical signals ^1^. Phenotypic heterogeneity, also referred to as extrinsic noise, occurs when cells respond differentially to the same external signal. Indeed, phenotypic heterogeneity is prevalent in multicellular systems, even when all the cells in the group share the same genetic background. Non-genetic heterogeneity, which may be due to stochasticity in gene expression or differences in gene expression, alternative splicing, and post translational modifications caused by external cues ^2–6^, may have severe implications in disease ^7, 8^ and developmental processes ^9–13^. However, it is not known whether non-genetic heterogeneity plays a constructive role in information transfer between cells in multicellular systems. As such, we ask whether some cells within a group function as leaders or followers, promoting the spread of information through the group, while others act individually, and whether such heterogeneity is important for the synchronization of multicellular dynamics.

Previously, we demonstrated that cell-cell communication through gap junctions modulated ATP-induced calcium signaling in monolayers of fibroblast cells ^14^. Tuning the levels of intercellular communications, by varying cell densities, inserting weakly-communicating cells, and by pharmacologically inhibiting gap junctions, controlled the temporal coordination of calcium signalling in neighboring cells ^14, 15^. To generalize these results and to elucidate how information transfer between single cells is integrated to synchronize population-level cellular responses, here we study collective mechanosensing in endothelial cells. Biologically, endothelial cells line the interior surface of blood vessels and form a monolayer that experiences varying levels of shear stress from blood flow ^16, 17^. Upon changing the flow rate (e.g., during acute wound), endothelial cells detect the change in shear stress, inform other cells such as smooth muscle cells, and adjust their internal signaling accordingly. Central to the cascade of events, shear stress leads to downstream ATP activation which modulates calcium signaling at the subcellular scale ^18–20^. As a group, the endothelial cells must coordinate their signaling dynamics to achieve a coherent and collective response. Specifically, intercellular calcium levels are synchronized via gap junction-mediated cell-cell communication ^14, 21^. Such synchronized calcium signaling is instrumental in modulating reepithelialization, angiogenesis, and extracellular matrix remodeling, which are essential processes in wound repair ^22–26^. Thus, we developed an integrated experimental-computational platform to quantitatively evaluate the roles that single cells take during the emergence of multicellular synchronization. Using this platform, we identified three key functions whereby single cells contribute to collective information processing that ultimately leads to multicellular synchronization. *Division of labor*, where single cells take differentiated functional roles in collective information processing; *Cell memory*, where single cells maintain and reinforce their specified functional roles in cell-cell communication in response to repeated external stimuli; And *information flow*, where the information gradually propagates spatially from the scale of single cells to eventually synchronize the collective.

## Results

### Endothelial cells in a monolayer synchronize their calcium dynamics in response to external shear stress

We employed a microfluidics system that can precisely control the temporal profile of the shear stress that the cells experience (Fig. 1A top). We grew confluent monolayers of HUVEC (Human Umbilical Vascular Endothelial Cell) cells on the bottom surface of the flow channels (Fig. 1A bottom). A computer interfaced flow switch regulated input pressure to induce smooth flow profiles in the microfluidic channel as verified by particle image velocimetry (Fig. 1B). The shear stress -induced calcium signal of the HUVEC cells was imaged with the fluorescent calcium indicator Calbryte-520 with single cell resolution (Fig. 1A inset, Video S1). We manually marked each cell center (Fig. 1A inset), recorded the intracellular calcium signal as a time-series of fluorescent intensity for every cell and verified that the magnitude of the cell’s calcium signal correlated with the magnitude of the corresponding flow shear stress that the cells experienced (Fig. 1C, Methods). This setting enabled us to investigate the collective calcium response of HUVEC cells when subjected to precisely controlled shear stresses.

**Figure 1:**
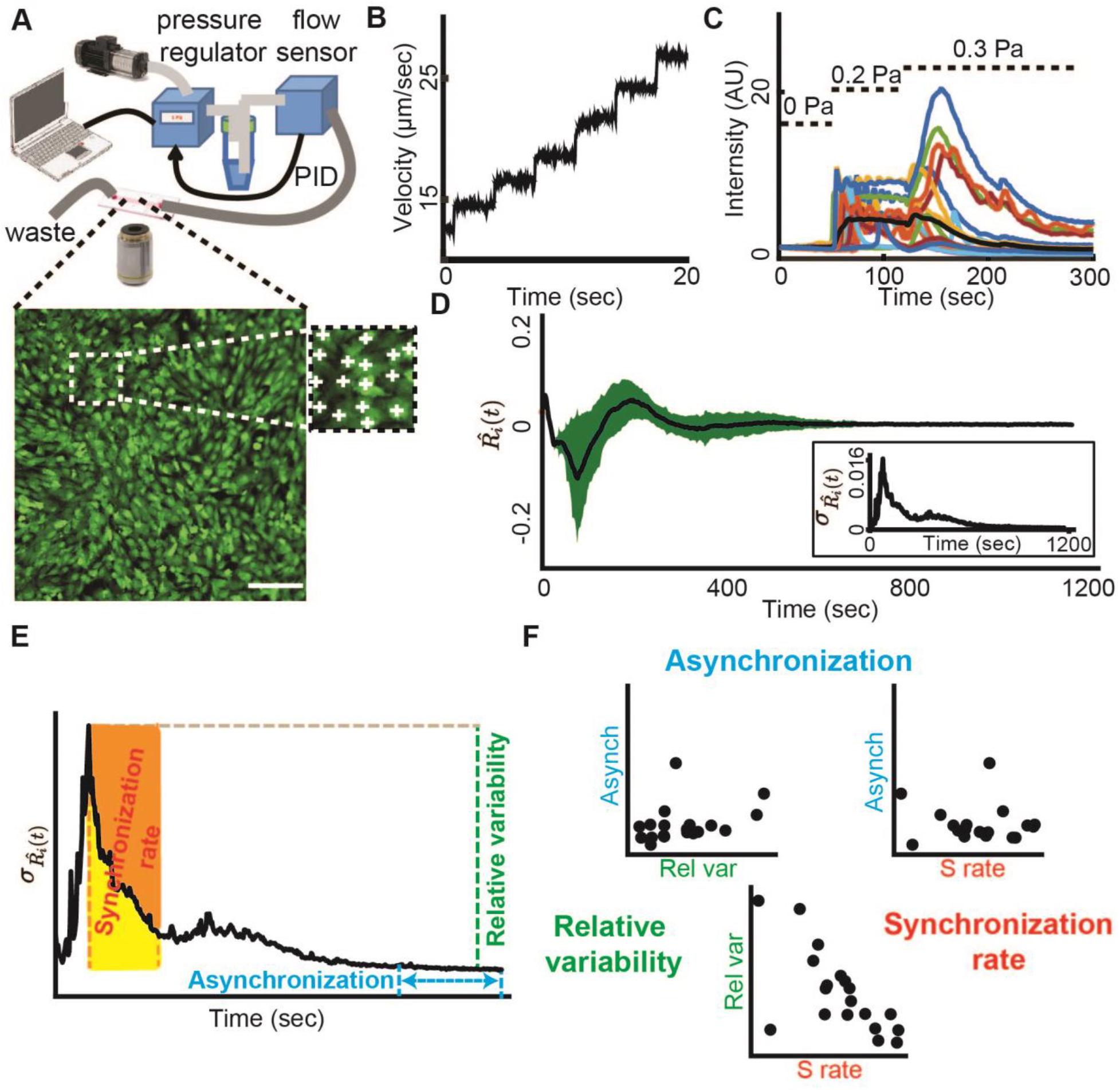
Collective calcium signaling of mechanosensing as a model to investigate the emergence of multicellular synchronization at the single cell resolution. (**A**) In a typical experiment, a monolayer of HUVEC was cultured in a microfluidic device where fluid flow applied shear stress on the cells. Top: Schematics of the setup. The input pressure that drives a laminar flow in the single channel microfluidics is controlled by a computer interface. The pressure is regulated in real time via a PID loop consisting of a pressure regulator and a flow sensor. Bottom: A monolayer of HUVEC loaded with the fluorescent calcium indicator Calbryte-520 as a readout of the cellular response to flow shear stress. Scale bar = 50 μm. Inset: manual annotation of single cells. (**B**) Particle image velocimetry verified that the regulated input pressure produces a smooth flow profile in the microfluidics channel. (**C**) Cells respond to step increase in the flow shear stress. Relative intensity is the relative change of the fluorescence intensity from the basal cell level (Methods). Dashed lines: individual cell calcium responses. Solid line: mean response of over 400 cells in the field of view. (**D**) Multicellular calcium dynamics is synchronized over time in response to external mechanical stimuli. The calcium dynamics of each cell was represented by the time-derivative of its relative fluorescent intensity 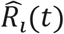. Black: mean calcium dynamics; Green: standard deviation. Inset: standard deviation of calcium dynamics over time. (**E**) Depiction of the three measures for synchronization overlayed on the plot of the standard deviation of calcium dynamics over time. Asynchronization (cyan) is the mean standard deviation over the final 200 seconds (100 frames) of the experiment. Relative variability (green) is the difference between the maximal and minimal standard deviation. Synchronization rate (red) is the ratio between the area under the curve of the collective asynchronization from the time of maximal variability in calcium dynamics for 200 seconds (100 frames, yellow area) and the area of the rectangle if no change in synchronization occurs during this time frame (yellow + orange area). See Methods for full description. (**F**) Correlation between synchronization measures. Relative variability versus synchronization rate Pearson correlation = −0.69, p-value = 0.0011. Asynchronization versus relative variability or versus synchronization rate Pearson correlation = 0.345 or −0.15 (both not statistically significant). Each observation represents a biological replica. N = 19 replicates (n = 6 experiments with pressure of 0.1 Pa, n = 13 with 0.2 Pa, see full distinction in Fig. S1).

Upon exposing the cells to a step-like increase in shear stress (0.1-0.2 Pa), which is similar to those that an endothelial cell experience during acute bleeding ^27^, the variability in the cells’ temporal derivative of their calcium signal (termed *calcium dynamics*, annotated 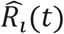 (Methods)) increased and then gradually reduced until it converged to a synchronized steady state (Fig. 1D). We defined three measures to quantify multicellular synchronization that relied on the population-level standard deviation 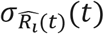 of single cell calcium dynamics 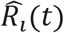 at different time points. First, *asynchronization* is defined as the time average of 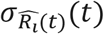 over the final 200 seconds. Low values of asynchronization implied improved synchronization across the entire group (Fig. 1E, marked in cyan). Second, 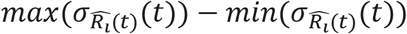, measures the magnitude of the transition from an unsynchronized to a synchronized state, which we termed *relative variability* (Methods, Fig. 1E, marked in green). Third, we defined the *synchronization rate* (Methods, Fig. 1E, marked in red) as the ratio between the area under the curve of 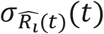 from the peak variability in calcium dynamics over 200 seconds (Fig. 1E, the area marked in yellow) with respect to the theoretical upper bound where the relative variability is zero (Fig. 1E, the combined areas marked in yellow and orange). We could not find a clear association between asynchronization and relative variability or synchronization rate, while the latter two measures were negatively correlated across experiments (Fig. 1F, Fig. S1). The mean time-correlation between neighboring cells, which we used previously ^15^, was associated with the multicellular synchronization but not associated with the relative variability or the synchronization rate (Fig. S2). These results demonstrate that multicellular synchronization is not necessarily correlated with the magnitude and speed of the response. We also found that increased levels of shear stress decreased multicellular asynchronization but not the relative variability or the synchronization rate (Fig. S3). Altogether, these results suggested that the cells formed a communication network that gradually evolved and reinforced synchronization despite the vast variability in single cell calcium response at the onset of shear stress application. In the following, we focus on relative low shear stress levels of 0.1-0.2 Pa to further elucidate the mechanisms of collective synchronization.

### Heterogeneity of single cell information transfer correlate with multicellular synchronization

We hypothesized that integrating and propagating information from the local scale, between single cells, to the global scale, drove the synchronization of an inherently heterogeneous multicellular network. To investigate this mechanism, we defined quantitative measures for cell-cell communications. If two cells communicate, we expect the past calcium dynamics of one cell to contain information regarding the future calcium dynamics of the other cell. Importantly, defined in this way, cell-cell communications can be bidirectional and asymmetric – cell A can influence its neighbor B differently from how cell B influences A (Fig. 2A). To quantify asymmetric cell-cell communication, we used Granger Causality ^28^ (GC), a classic statistical method from the field of information theory, to infer cause-effect relationships between cell pairs from their fluctuating calcium dynamics. Granger Causality uses linear regression to quantify the extent to which the prediction of values in one time series can be improved by including information from another time series. This provides us with an established framework to extract feedback and feedforward relations from pairwise variables’ fluctuating time-series.

**Figure 2:**
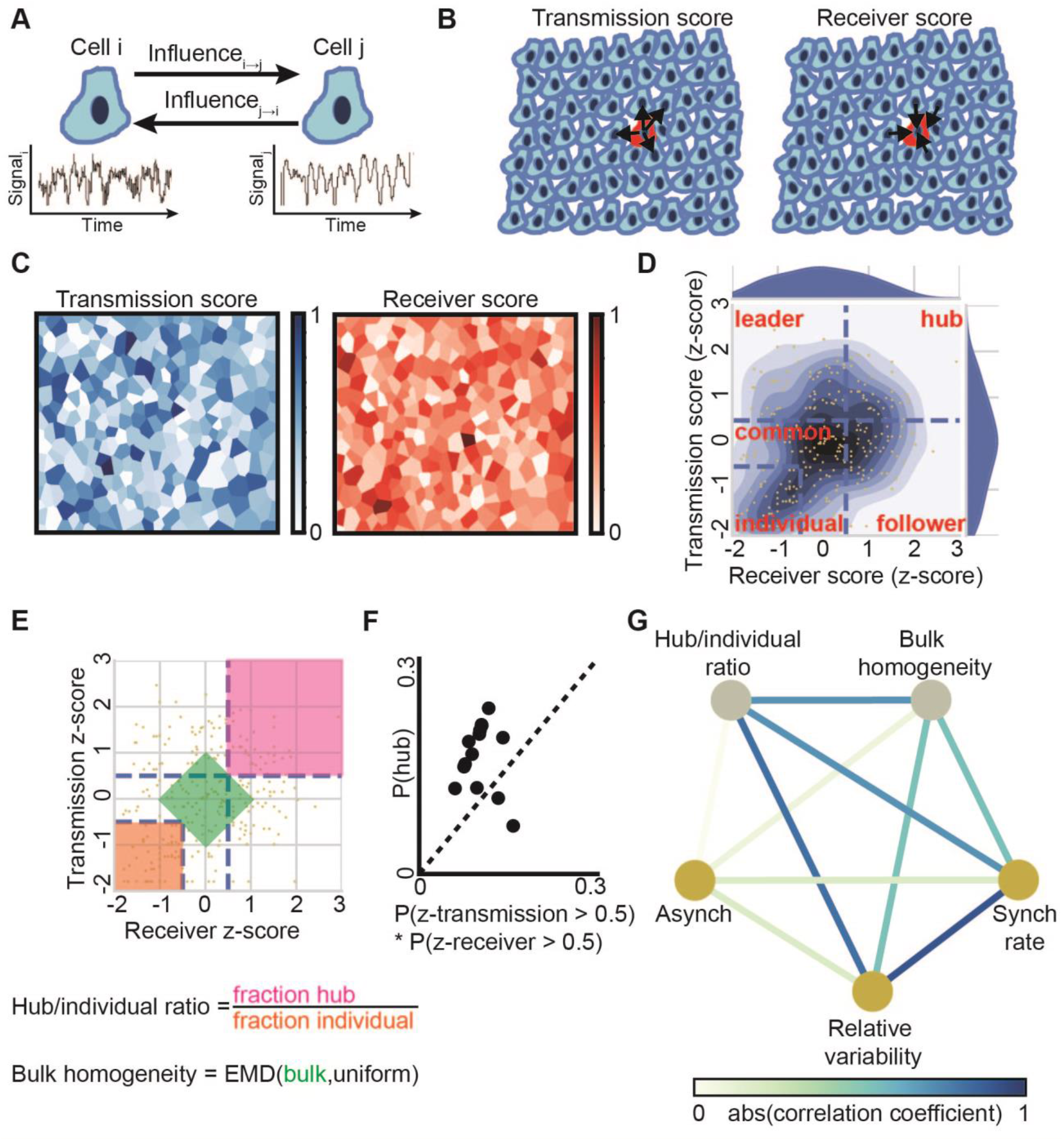
Correlating single cell information transfer and multicellular synchronization. (**A**) Schematics of cell-cell communication. Generic estimation of the asymmetric mutual influence between a pair of cells from their fluctuating time series. The influence of cell i on cell j is defined as the extent to which the past signal of cell i improves the prediction of cell j’s signal beyond the past signal of j alone and is determined using the pairwise asymmetric Granger Causality statistical test. (**B**) The transmission and the receiver scores are calculated as the probability for a significant outgoing (respectively, ingoing) Granger Causality at topological distance of up to two (nearest and next-to-nearest neighbor cells). (**C**) Visualization of the spatial single cell heterogeneity of the transmission/receiver score. (**D**) Kernel Density Estimate plot visualization of the normalized transmission and receiver score (blue gradient contours). Partitioning of the z-score normalized transmission-receiver space to five regions (blue dashed lines), their corresponding annotations or “roles” (red text) and single cell assignment (yellow dots). Individual: transmission and receiver z-score < −0.5, Common: transmission z-score in the range of −0.5-0.5 and receiver z-score < 0.5 or receiver z-score in the range of −0.5-0.5 and transmission z-score < 0.5. Leader: transmission z-score > 0.5 and receiver z-score < 0.5. Follower: receiver z-score > 0.5 and transmission z-score < 0.5. Leader: transmission and receiver z-score > 0.5. (**E**) Depiction of the two measures for heterogeneity overlayed on the normalized transmission and receiver score plot. Hub/individual ratio - the ratio between the fraction of cells in the communication hub role (magenta) and the fraction of cells in the individual role (orange). Bulk homogeneity - the distance (earth movers distance, EMD) between the experimentally observed and the uniform distribution for all cells that hold |transmission score + receiver score| < 1 (green). (**F**) Communication hubs were enriched beyond their expected probability. p-value = 0.0076 via Wilcoxon rank-sign test on P(hub)-P(z-transmission)*P(z-receiver). For panels F and G, N = 14 biological replicates at 0.1 or 0.2 Pa (see S7D). (**G**) Pairwise associations between two heterogeneity measures (in gray: hub/individual ratio, bulk homogeneity) and three synchronization measures (in yellow: asynchronization, relative variability, synchronization rate). Edges color represents the level of association, as quantified by magnitude of correlation coefficients, between a pair of measures. Note that some of the measures are positive measures for synchronization properties (relative variability, synchronization rate) and heterogeneity (hub/individual ratio), while other properties are negative measures (asynchronization and bulk homogeneity) so negative correlations between measures from these two groups imply associations between synchronization and heterogeneity (for example, the negative correlation between bulk homogeneity and synchronization rate). N = 14 biological replicates per association. Full data in Fig. S7, and Fig. S9.

To avoid spurious cause-effect relations, Granger Causality requires the time-series being analyzed to be stationary, i.e., fluctuating signals with a consistent mean and variability. Therefore, we excluded experiments where less than 85% of the cells passed two stationary tests (Kwiatkowski–Phillips–Schmidt–Shin ^29^ and Augmented Dickey–Fuller ^30^, Fig. S4, Methods) and in the remaining 14 (out of 19) experiments we analyzed only the cells that passed both stationary tests. When one cell’s calcium dynamics significantly contributed to the accurate prediction of another cell’s signal we defined a directed GC edge from the first cell to the other (Methods). For every cell in the monolayer we calculated the *transmission score* and the *receiver score* as measures for being influential or influenced-by cells in its local vicinity -- nearest neighbor cells and next-to-nearest neighbor cells, i.e., cells with topological distance up to 2. We defined the transmission score as the probability of outgoing GC edges, and the receiver score as the probability of ingoing GC edges (Fig. 2B) (Methods). Cells took different roles in the multicellular communication network as indicated by the spatial heterogeneity in their transmission/receiver scores (Fig. 2C), which was not associated with the number of cell neighbours (Fig. S5). We normalized the single cell transmission and receiver scores across the population by calculating for each cell its receiver and transmission z-score - the number of standard deviations away from the mean (Fig. 2D, Methods). The normalized scores allowed direct comparison of the single cell heterogeneity across experiments.

To identify relations between heterogeneous single cell communication properties and multicellular synchronization, we partitioned the normalized transmission-receiver space into five regions and assigned the cells to groups according to the region they occupied. *Individual cells*, whose calcium dynamics were independent of cells in their local vicinity, have normalized transmission and receiver scores both below 0.5. *Common cells*, with average communication properties. *Leader cells*, with high transmission scores (transmission score > 0.5 and receiver score < 0.5), *follower cells*, with high receiver score (receiver score > 0.5 and transmission score < 0.5), and *communication hub cells*, characterized by both transmission and receiver normalized scores above 0.5 (Fig. 2D, Methods). This data-driven partitioning defined five distinct roles that cells take in terms of information transfer in the multicellular communication network. We correlated the fraction of cells at each role to the synchronization measures but did not find an association between any of the 5 x 3 combinations of fraction of cells at a specific role and a synchronization measure correspondingly (Fig. S6). Strikingly, we discovered a remarkable association between the ratio of the fraction of communication hub and of individual cells, termed *hub/individual ratio* (Fig. 2E), to the relative variability and to the synchronization rate, but not to the asynchronization (Fig. S7A-C). These results suggest that increased fraction of communication hub cells along with decreased fraction of individuals were associated with an improved synchronization process (reduced relative variability and increased synchronization rate). Importantly, the hub/individual ratio is a local cell property - dependent on the spatial organization of cells in the vicinity of cells playing the hub or individual phenotype, as verified with a spatial permutation test (Fig. S8A). Thus, these results established a link between the “local” single cell communication properties and the process leading to the “global” emergent property of multicellular synchronization.

We next focused our attention to the fraction of cells taking the “communication hub” role. The assignment of a cell to the “hub” role was determined as 0.5 standard deviations above the mean in both the receiver and transmission score. Assuming that the transmission and receiver scores of a cell are independent (or negatively correlated), the expected fraction of “hubs” in the population would be bounded by P(zscore_transmission > 0.5) * P(zscore_receiver > 0.5). However, the observed fraction of cells in the communication hub role was enriched beyond this expectation across experiments (Fig. 2F, Fig. S7D).

The hub/individual ratio is a measure for heterogeneity, relying heavily on the extreme high and low values in the transmission and receiver scores distributions (Fig. 2E, top right and bottom left corners). More hubs increase this ratio, and more individuals decrease it. But how do we quantify the heterogeneity at the bulk of the transmission/receiver score space? We defined a second measure for heterogeneity of all cells with |transmission score + receiver score| < 1, which consists of approximately 40% of the cells in the population (Fig. 2E, central region). We calculated the distribution of the absolute sum of the transmission and receiver scores for the cells in this bulk region and defined the *bulk homogeneity* as a scalar value measuring the deviation from the uniform distribution (Methods, Fig. 2E). For high heterogeneity experiments, the bulk homogeneity was low, following a uniform distribution, while in more homogeneous experiments, the bulk homogeneity was higher as the data distribution deviated from uniformity. We used the Earth Mover’s Distance (EMD) ^31, 32^ to calculate dissimilarities between the observed and the uniform distributions. The EMD of 1-dimensional distributions is defined as the minimal ‘cost’ to transform one distribution into the other ^33^. Larger values reflect a more homogeneous distribution of the population’s bulk single cell communication properties. The cells considered for the bulk homogeneity calculation did not include hubs or individuals and thus defined an independent and complementary measure for heterogeneity. Still, bulk homogeneity was correlated with the hub/individual ratio (Fig. S9A). Bulk homogeneity was verified to be a local cell property that depends on the spatial organization of cells in the vicinity (Fig. S8D-E). Bulk homogeneity was associated with the relative variability and with the synchronization rate, but not to the asynchronization (Fig. 2G, Fig. S9B-D).

Together, these results suggest that the heterogeneity in the roles that single cells took in the multicellular network, or “division of labor”, was associated with the process leading to the emergence of collective calcium synchronization, where “communication hubs” were enriched. As a note, none of the heterogeneity or the other synchronization measures was associated with the asynchronization, which was not even significantly associated with the mean *GC edge probability* (Fig. S10).

### Experiments with periodic mechanical stimuli reveal that information flow is associated with multicellular calcium synchronization

After characterizing the communication networks exhibited by endothelial cell monolayers to shear stress, we asked if the network could be trained to adapt to changing external stimuli. To this end, we extended our assay to multiple rounds of repeated mechanical stimuli (Video S2). By treating each round as an independent *cycle*, and comparing single cell responses across cycles, we could focus on the evolution of synchronization in the multicellular communication network (Fig. 3A, Methods). We found that the HUVEC monolayer gradually reinforced synchronization as observed by the gradual decrease in the standard deviation of the cells’ calcium dynamics 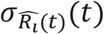 (Fig. 3B). This trend of improved synchronization coincided with a gradual increase of the cell’s mean GC edge probability (Fig. 3C) and the mean receiver and transmission scores across the population along the progression in cycles over time (Fig. 3D-E, Fig. S11, Video S3, Video S4, Video S5). These results established that multicellular synchronization was associated with the GC edge probability, a proxy for the overall information flow within the multicellular network.

**Figure 3:**
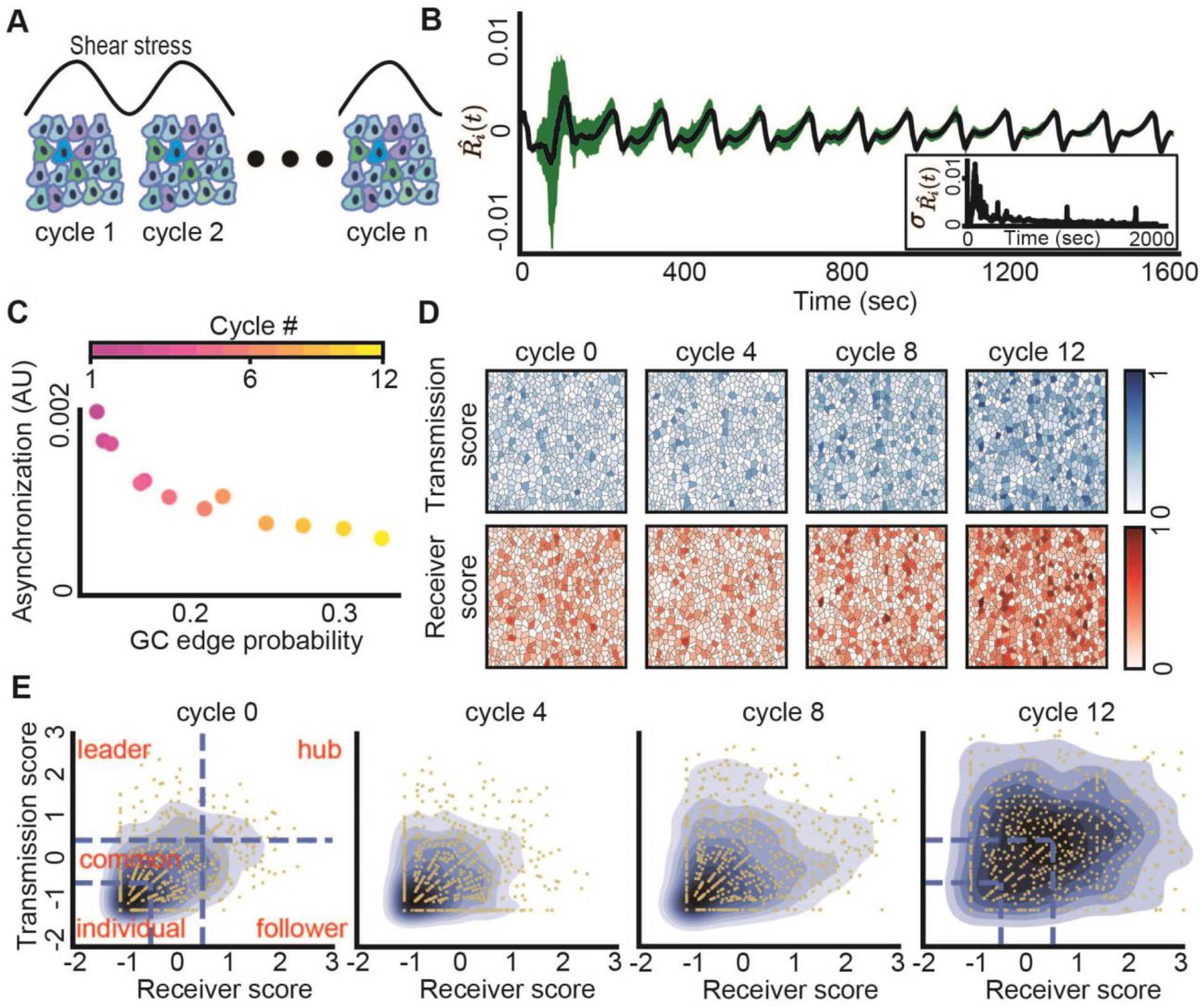
Information flow is associated with multicellular synchronization. **(A)** Depiction of the periodic mechanical stimuli experiment setup that included 13 cycles of continuous shear stress in 6 biological replicates (Methods). **(B)** Multicellular calcium dynamics is synchronized over time in response to periodic external continuous mechanical stimuli. Each peak represents a middle time point of the single cycle. In total there are 13 cycles. Black: mean calcium dynamics; Green: standard deviation. Inset: standard deviation of calcium dynamics over time. **(C)** Asynchronization was negatively associated with GC edge probability. Pearson’s correlation = −0.8933, p-value < 0.0001. Colormap represents the cycles (time), the multicellular network became more synchronized with increased GC edge probability as the cycles progressed. This analysis considered 12 cycles. The first cycle was an outlier and was excluded from this analysis. **(D-E)** The transmission and receiver scores evolution over the cycles. Shown are all cells in cycles 0, 4, 8, and 12. (**D**) The transmission and receiver scores increased over the cycles. Shown are the cells color coded according to their transmission (top, blue) and receiver (bottom, red) scores. **(E)** Kernel density estimate plot visualization of the normalized transmission and receiver score over the cycles (blue gradient contours). Left: partitioning of the z-score normalized transmission-receiver space to five regions (blue dashed lines), each cell (yellow dot) was assigned to a group or “role” (red text) according to the region they resided at.

### Functional cell memory: cells maintain their roles in the communication network and reinforce them over time

We next asked to what extent the communication properties of cells were intrinsic cellular properties. To this end we correlated single cells’ transmission and receiver scores across the repeated mechanical stimulus cycles while testing the null hypothesis that these scores were assigned randomly between consecutive cycles. We found that single cells’ transmission and receiver scores were strongly correlated between consecutive stimulus cycles, that this correlation gradually increased as cells underwent additional stimulus cycles (Fig. 4A, Fig. S12A), and could not be explained by the autonomous cells’ response to the external mechanical stimuli (Fig. S13). Measuring single cell correlation between larger temporal gaps of 2-4 cycles did not show a dramatic diminishing pattern suggesting that the cellular memory is stable for time scales of at least 4-8 minutes (Fig. S12B). These results suggest that cells maintain and gradually reinforce memory regarding their role in the multicellular communication network at the timescales relevant for collective synchronization.

**Figure 4:**
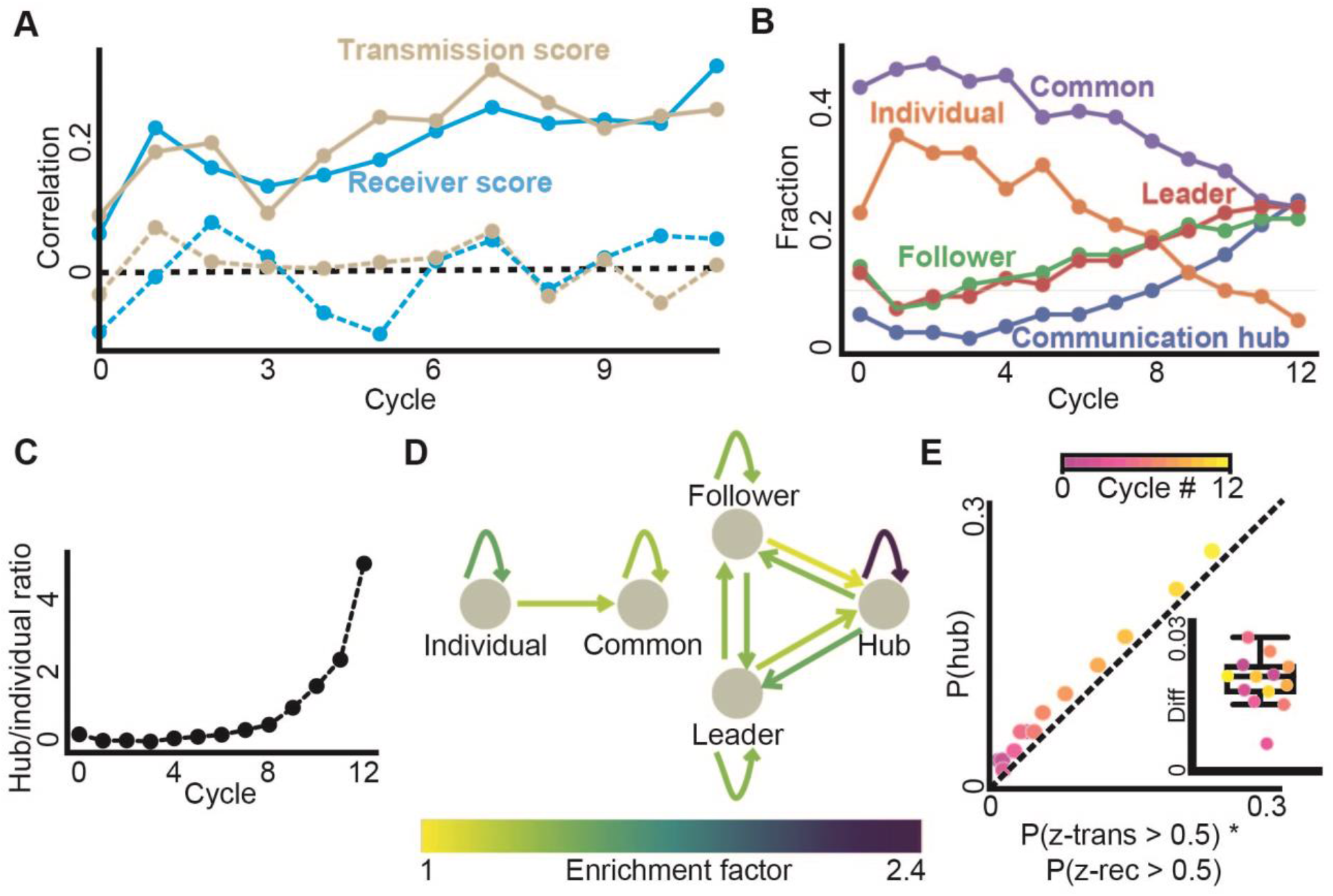
Cells’ memory and state transitions. (**A**) Cells transmission and receiver scores are correlated across consecutive cycles (solid lines), are reinforced over time (Pearson coefficient = 0.7512, p < 0.0001), and are a local cell property as validated with permutation analysis - shuffling the cells in the next cycle and calculating correlation (dashed line, see Methods). P-value for the significance of the memory ≤ 0.001 (except the first cycle: p-value of transmission and receiver score 0.021 and 0.15 correspondingly, and the third cycle’s transmission score p-value of 0.017, Fig. S12A). (**B**) Fraction of cells at each communication role over the cycles. (**C**) Hub/individual ratio over the cycles. (**D**) Enrichment factors of cellular state transition. Depiction of the single cell transitions between functional roles that were enriched beyond the expected values of a null model. The null model was based on the marginal distribution of the roles (Fig. S14, Methods). Shown are edges with fold increase over 1 (color code). (**E**) Enrichment in communication hubs over cycles (color code). Expected fraction of communication hub cells based on the fraction of cells with transmission score and receiver scores above 0.5 (x-axis) versus the actual fraction of communication hubs (y-axis), values above the Y = X line indicate enrichment in communication hubs. Inset shows the signed difference between the observed (P(hub)) and the expected (P(z-trans > 0.5) * P(z-rec > 0.5)), where all cycles show positive values, N = 13, Wilcoxon sign-rank test p-value = 0.0015.

The combined effect of the increasing information flow and the functional cell memory was associated with a gradual increase in the fraction of followers, leaders and communication hubs and a decreased fraction of common and individual cells (Fig. 4B, Video S6), leading to an increased ratio between “hub” and “individual” cells that was correlated with the increased synchronization (Fig. 4C). To follow the dynamic trajectory of single cells between roles we analyzed cell state plasticity, the probability of transitioning from one state (functional role) to another cell state in consecutive cycles. In particular, we computed the enrichment factor -- transition probabilities between any two states and normalized the quantity by the fully random transition probabilities (Methods, Fig. S14). As expected from our earlier observation of functional memory (Fig. 4A), we found that cells tended to maintain their roles or “similar” roles, as reflected by self-transition enrichment factors above one (Fig. S14, Fig. 4D). Generally, single cells followed a temporal trajectory from the roles characterized with less communication capacity to roles with increased communication (Fig. 4D - showing edges only for enrichment factors > 1). Intriguingly, we found symmetric transition folds between the follower-and leader roles, and the transition from communication hub to the follower/leader roles was enriched compared to the opposite transition to a communication hub (Fig. 4D, Fig. S14). The communication hub role was found to be much more stable than other roles or transitions (2.4 fold dwell probability compared with a fully random process), explaining their rapid spread in the population (Fig. 4B). The fraction of communication hubs also systematically exceeded its expected fraction if transmission and receiver scores were independent. Along with the gradually increasing information flow, the results highlight the role of functional memory in communication hub enriching (Fig. 4E).

### Information gradually propagates from the (local) single cell to the (global) multicellular scale

Our data associates the heterogeneity of cells’ transmission and receiver scores, memory (and reinforcement) of the single cell’s communication properties and information flow with multicellular synchronization. To test our hypothesis that the synchronization process is driven by effectively propagating information from the local scale (between single cells), to the global (collective) scale, we measured to what extent local cell properties explained the information flow in the multicellular network. First, we computed the neighboring pair cross correlation coefficients for direct observations and spatially permuted data. We found that the spatial permutation always decreased the cross correlation, therefore cross correlation was maintained as a local cell property throughout the experiment even in the presence of common external stimuli (Fig. 5A, upper-left inset). Intriguingly, spatial permutation decreased the GC edge probability in early cycles but increased the edge probabilities in later cycles. (Fig. 5A main panel and the lower-right inset). These results indicate that once neighboring cells reach sufficient synchronization their ability to influence each other is less effective than cell pairs far apart. We validated these observations more systematically by correlating the topological distance between pairs of cells to their GC edge probability (Methods). This analysis established that at the onset of the experiment, the information flow is dominated by local cell-cell interactions and is gradually transitioning to the global scale as the multicellular network synchronizes (Fig. 5B).

**Figure 5:**
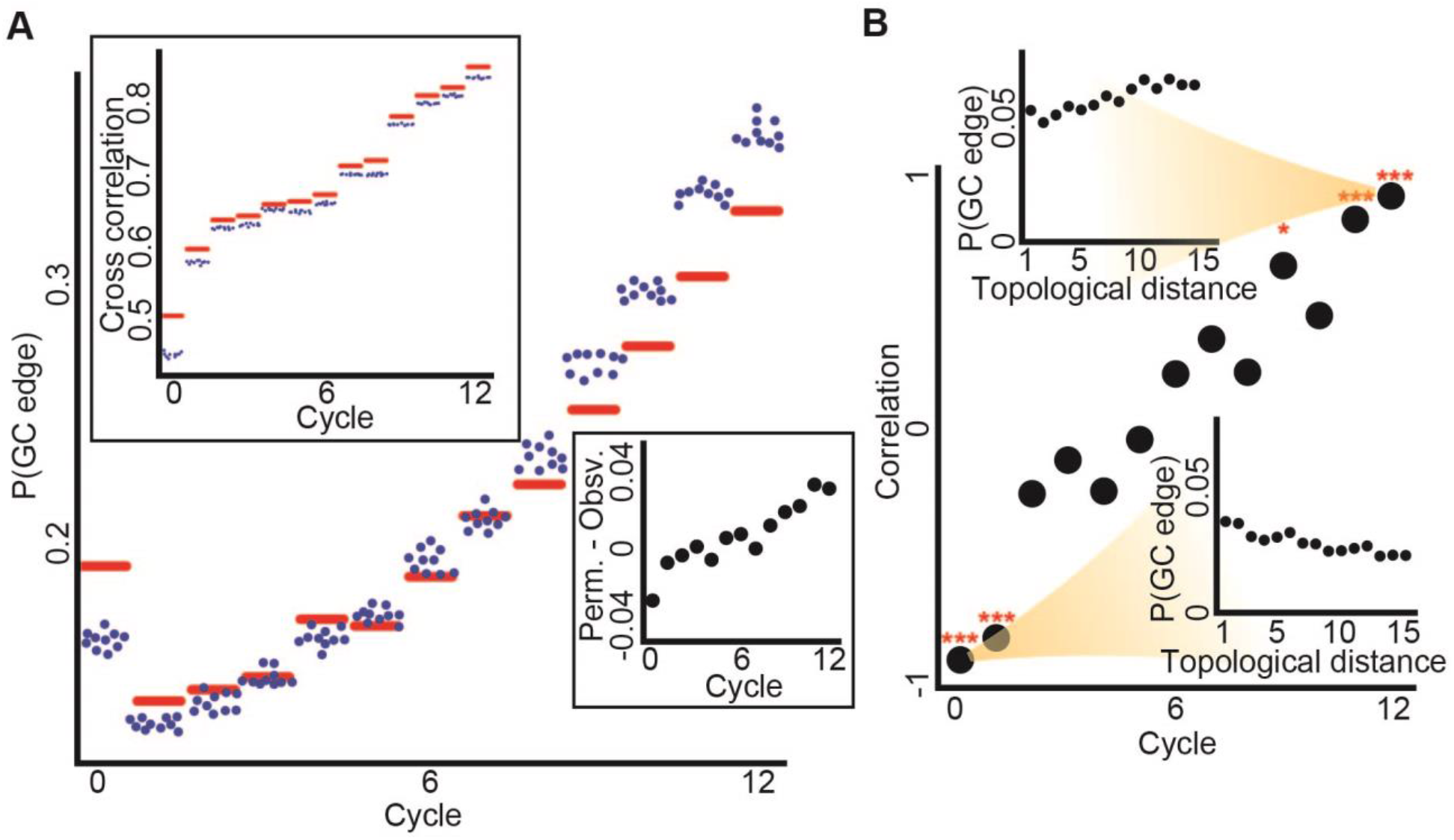
Gradual local to global transition in information spreading. (**A**) Main panel: the observed versus permuted Granger causality edge probability, P(GC edge), over the cycles. Upper left panel: the mean observed versus mean permuted neighbor cross correlation over the cycles. For both panels the red horizontal line is the experimental observation, while each blue dot is the result of one of ten independent spatial cell permutations. Bottom right inset: experimental GC edge probability subtracted from the mean permuted GC edge probability using the same data as in the main panel. Through cycles 0 to 12 Pearson coefficient = 0.94, p-value < 0.0001. (**B**) In the main panel each dot represents the Pearson correlation between the topological distances of pairs of cells to the corresponding GC edge probability in a given cycle. Through cycles 0 to 12 Pearson coefficient = 0.964, p-value < 0.0001. *** - p-value < 0.0001, * - p-value < 0.05, for Pearson correlation significance test. Insets show the P(GC edge) as a function of the topological distance between cell pairs for the first (bottom right) and last (top left) cycles. For this analysis, we randomly selected for each cell at each topological distance at most ten neighboring cells due to computational cost and performed FDR multiple hypotheses correction (see Methods for full details).

## Discussion

The emergence of robust multicellular behaviors from heterogeneous single cell dynamics is a poorly understood, but fundamentally important phenomenon in living systems ^34^. Here we provide insights in bridging the scales between local cell-cell communication and global multicellular synchronization. This was achieved by measuring asymmetric information transfer at single cell resolution in multicellular monolayers under externally applied mechanical stimuli. By employing Granger Causality to systematically quantify the communication of a cell with other cells in their local environment, we defined for each cell its capacity to transmit and to receive information in the multicellular communication network. Our method relies on local pairwise analysis of cell dynamics, and does not require explicitly constructing the network or committing to a specific network architecture. This model-free data-driven approach can be applied to a broad set of biological systems from synchronized beating of cardiomyocytes ^35^, intercellular communication through the microenvironment ^36^, brain activity ^37^, molecular signaling ^38^ and coordinated cell migration ^39^.

We showed the cells were actively communicating with one another locally, as supported by multiple lines of evidence throughout our study. First, we demonstrated that both hub/individual ratio and bulk homogeneity depended on the spatial organization of cells in their vicinity (Fig. S8). Second, we found that the activation time, a cell’s autonomous response to the external stress, was not associated with the transmission or receiver score (Fig. S13A), which would also be conflicting with the enrichment of communication hubs (which are both leaders and followers). Third, cells “remembered” and reinforced their roles in the multicellular communication network over time, as a local, spatially-dependent property (Fig. 4A), but did not “remember” their activation time in previous cycles (Fig. S13B). Fourth, neighbor pair cross correlation was a local cell property throughout the experiment (Fig. 5A). Together, our data established the decoupling of the local cell-cell communication from the global external stimuli, and excluded the possibility that the emergence of multicellular synchronization was an artifact caused by differential cell autonomous response to the external shear stress that was applied on the group.

Our data reveal that single cells take different roles in cell-cell communication (heterogeneity or “division of labor”), gradually reinforce these roles (“functional cell memory”), and increase the connectivity (“information flow”) in the multicellular network. These three mechanisms work in concert to synergistically contribute to the emergence of multicellular synchronization. We hypothesize that cell heterogeneity expands the dynamic range of multicellular responses that cell memory stabilizes the dynamics against intrinsic and extrinsic noise and that information flow sustains and reinforces the multicellular dynamics.

We found that heterogeneity in cells’ communication properties were associated with improved convergence to synchronization (Fig. 2G). We also observed that the fractions of cells at each functional role, excluding individuals, became more balanced through periodic cycles (Fig. 3E and 4B), in agreement with our hypothesis that heterogeneity constructively contributes to the synchronization of a noisy multicellular system. Cell heterogeneity could arise from stochastic gene expression levels, signaling kinetics, physiological cellular states such as cell cycle, and/or microenvironmental cues ^6, 40–45^. The rapid time scale of the systematic changes in single cell communication properties suggests that additional mechanisms may be involved such as cross-talk between mechanosensing and gap junctional communication. Consistently, the network evolution in multi-cycle experiments also suggested that rapid modulations such as through kinetics of calcium signaling, contributed to the observed heterogeneity.

Previous studies have reported multiple sources of microenvironment-dependent cell memory. For instance, cells can remember past mechanical properties of their substrate, which influence their differentiation ^46^. Cells can also sense changes in their extracellular signal by remembering past extracellular stimulation via a receptor-mediated mechanism ^47^. In the context of collective cell migration, a recent study showed that cells remembered their polarized state independently of cell–cell junctions ^48^, and another study revealed associative memory of electric field and chemoattractant at stimuli in a unicellular organism migration patterns ^49^. In our study, single cell memory of communication properties contributes to the temporal evolution of the multicellular network to its synchronized state. The dissociation between a cell’s activation time and its functional role in information processing underpins the dynamic nature of memory, which is also consistent with the unidirectional evolution of the multicellular network (Figs. 4–5).

Our study reveals a self-organized multicellular network that supports information flow from local to global scales. Such information may be carried by two main signaling mechanisms, juxtacrine (contact-dependent) and autocrine (secreted-dependent) ^50^. A juxtacrine channel allows a cell to establish conversation with its (physically touching) immediate neighbors without interference from extracellular space. For HUVEC cells such communication can be realized by gap junctions ^51^. On the other hand, an autocrine channel allows a cell to broadcast its information through diffusive messengers in the extracellular space. For HUVEC cells stress-triggered ATP release and ATP-induced calcium dynamics constitute an autocrine pathway ^52^. While both mechanisms could contribute to the information flow within the multicellular network, we suggest contact-dependent signaling as the dominant mechanism. While a recent study suggested that positive feedback of a diffusive signaling mechanism can drive accelerated, long-range information transmission ^53^, the external flow in our system is likely to rapidly dilute the diffusive messenger ^54^. The contact-dependent information flow hypothesis is also supported by our previous studies where we demonstrated that blocking gap junctions, or inserting weakly communicating cells impared the information flow ^14, 15^.

Altogether, our results suggest the following phenomenological model for multicellular synchronization. Cells are gradually “learning” the local network structure around them (heterogeneity), adjusting their internal state, reinforcing it (memory), and thus stabilizing the network architecture. This stabilized network structure reduces conflicting communication interferences and thus promotes enhanced spread of information from the local to the global scale to eventually synchronize the group.

## Methods

### Cell culture

Human Umbilical Vascular Endothelial Cells (HUVEC) were purchased from Lonza and were cultured following the vendor’s instructions. To prepare samples, cells were detached from culture dishesusing TrypLE Select (Life Technologies) and suspended in growth mediums before being pipetted into the microfluidics devices at cell density of approximately 1000 cells/mm2 allowing the cells to form monolayers. After overnight incubation, fluorescent calcium dye (Calbryte 520, AAT Bioquest) was loaded for 40 minutes prior to imaging.

### Microfluidics

The organic elastomer polydimethylsiloxane (PDMS, Sylgard 184, Dow-Corning) used to create the microfluidic devices was comprised of a two part mixture - a base and curing agent - that were mixed in a 10:1 ratio, degassed, and poured over a stainless steel mold before curing at 65°C overnight. Once cured, the microfluidic devices were cut from the mold, inlet/outlet holes were punched, and the device was affixed to a No. 1.5 coverslip via corona treatment. The cross section of the flow chambers was rectangular (2 mm X 1 mm). See Fig. 1A for depiction.

### Applying controlled shear stress on the cells

The microfluidics flow rate was controlled by a PID-regulated pressure pump and was monitored using an inline flow sensor (Elveflow). To verify the stability of the flow profile we mixed 1 micrometer fluorescent particles in the solution and used particle image velocimetry to quantify the flow rate (Fig. 1B). To calculate the shear stress, we approximated the flow profile in the flow chamber as low-reynolds number pipe flow. We considered the cells in the field of view to experience uniform shear stress calculated at the center of the flow chamber. This was possible because the imaging window was narrow (470 μm) compared with the chamber width (1 mm).

In the “step” experiments we exposed the cells to a “step”-like shear stress of 0.1, 0.2, 0.6, 1 or 1.6 Pa for approximately 20 minutes. In the “cycles” experiments we applied multiple rounds of 2 minute long global external periodic mechanical stimuli.

### Live cell imaging

We imaged the calcium dynamics of HUVEC cells using a 20X magnification oil immersion objective lens (for step-stress experiments) and a 10X dry lens (for cyclic-stress experiments, to avoid flow-induced focus drift). The fluorescent images were captured with a CMOS camera (Hamamatsu Flash 2.8) at 0.5 Hz and 1 sec exposure time. The images were stored as tif files of 960 × 720 pixels with physical pixel size of 0.65 μm X 0.65 μm (under 20X) or 1.2 μm X 1.2 μm (under 10X).

### Measuring single cell calcium signaling

We manually annotated every cell center (Fig. 1A inset), and recorded the mean fluorescent intensity of approximately 40 μm^2^ around the cell’s center as a proxy of the intracellular calcium signal time-series of each cell. We normalized the calcium signal according to the approach described in ^55^ (“response curve normalization”). The response curve of each cell was defined as 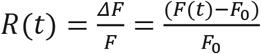, where *F*(*t*) was calculated as the mean fluorescent intensity at time t, The baseline *F*_0_ was calculated as the mean of the first 5 frames (10 seconds) of *F*(*t*) before the mechanical stimulation was turned on. Temporal long-pass filter smoothes the R(t) time series to reduce the effects of outliers. Temporal smoothing was performed using Python’s lowess function from statsmodels (statsmodels.nonparametric.smoothers_lowess) with the parameter frac=0.01. R(t) provides us with a dimensionless measure for the intracellular calcium magnitude relative to the cell’s basal intensity.

To evaluate the change in the calcium signal in response to external mechanical stimulus we derived the cells’ calcium signal in time: 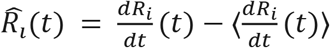, where 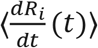 is the mean value of 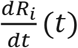, and 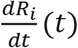 was calculated by Python’s numpy convolution operator over the time series with the parameter values mode=’same’ and with a five point stencil filter [1, −8, 8, − 1] where the result was divided by constant of 12. A cell’s calcium signal temporal derivative 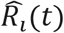 was termed calcium dynamics.

### Measuring synchronization

We defined three measures to quantify multicellular synchronization that relied on the standard deviation of single cell calcium dynamics at different time points 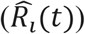.

The first measure was *asynchronization*, which measured the mean standard deviation of the cells’ calcium dynamic at the cells’ synchronized state (Fig. 1E, marked in cyan). Here we defined 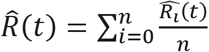, where *n* is the number of cells, as the mean calcium dynamic of all the cells at time *t* and 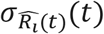 function as the standard deviation of the cells’ calcium dynamic. The asynchronization was calculated as the mean 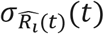 of the final 200 seconds. In the cycles experiment, the mean 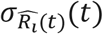 was calculated over the entire cycle time. Low values implied improved synchronization across the entire group.

The second measure was *relative variability* (Fig. 1E, marked in green), which measured the magnitude of the transition from an unsynchronized to a synchronized state by calculating 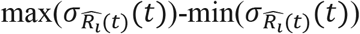.

The third measure was *synchronization rate* (Fig. 1E, marked in red), which measured the ratio between the area under the curve of 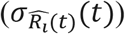 from the peak variability in calcium dynamics over 200 seconds (Fig. 1E, the area marked in yellow) with respect to the theoretical upper bound where the relative variability is zero (Fig. 1E, the combined areas marked in yellow and orange). The area under the curve was calculated using trapezoidal rule where the space between sample points is 2 sec (1 frame), 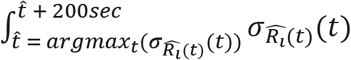.

### Granger causality

Granger causality (GC) is a statistical method to quantify the information flow among multiple variables’ time-series ^28^. Intuitively, time-series B is said to be “Granger causal” of time-series A, if the variability of A can be better explained by previous values of B and A, compared to using only previous values of A. Granger causality is an approximation to “transfer entropy” and under the assumption of Gaussian distribution it is exactly equivalent ^56^.

Formally, given two-time series *x_i_*(*t*) and *x_j_*(*t*), where t∈Z. The autoregressive model of *x_i_* is:

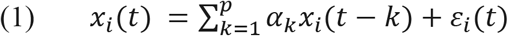

Where, p is the lag order, the number of previous observations used for prediction, *α_k_* is the coefficient of *x_i_* and *ε_i_* is the prediction error at time t. The autoregressive model of *x_i_* based also on the previous observations of *x_j_* is:

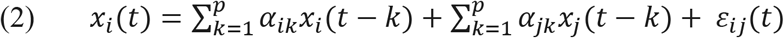

Where, p is the lag order, *α_ik_* is the coefficient of *x_i_*(*t* – *k*), *α_jk_* is the coefficient of *x_j_*(*t* – *k*) and *ε_ij_*(*t*) is the joint error of *x_i_* and *x_j_* predicting *x_i_*.

### Stationarity test

To avoid spurious causality connection, *x_i_* and *x_j_* both must comply with a stationary process before applying the granger causality test. Intuitively, stationary means that the statistical characteristics such as average and variance of a time series are independent of time. For each cell’s 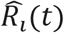 time series we applied two statistical tests for stationarity. First, Kwiatkowski–Phillips–Schmidt–Shin (KPSS) ^29^ tests the null hypothesis of stationarity against the alternative of unit root. Second, Augmented Dickey–Fuller (ADF) ^30^ was applied as a complementary test for KPSS and tests the null hypothesis for unit root against the alternative of stationary. We excluded five of the nineteen “step” experiments where less than 85% of the cells passed both the KPSS and the ADF stationary tests with significance of below 0.05 (Fig. S4). From the remaining experiments we considered only cells with time-series that passed both stationarity statistical tests.

### Pairwise calibration of the lag order

Granger Causality is based on linear regression and thus sensitive to the lag order, i.e., the number of past time frames used to make future predictions. In the context of a time-series, the autoregressive (AR) model is the estimator of the next time point value based on its own previous values. Higher lag-order reduces the bias but increases the variance while lower lag-order reduces the variance but increases the bias ^57^. We selected the lag order, for each cell pair independently, as the minimal lag derived from four methods: Akaike information criterion ^58^, Bayesian information criterion ^59^, Final Prediction Error ^60^ and Hannan–Quinn information criterion ^61^. The minimum lag was selected to avoid overfitting without losing information backup by using Portmanteau test which checks for whiteness (i.e, the error does not contain a pattern) ^62^.

### Granger causality statistical test

We applied a statistical test to infer granger causality between two 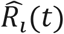 time series *x_i_* and *x_j_* (denoted, *GC_x_j_ −>_x_i___*) ^28^. GC tests the null hypothesis that *x_j_* is not contributing to the explained variance of *x_i_* (Equation (2)) in relation to the model derived solely from past values of *x_i_* (Equation (1)). This null hypothesis is rejected when at least one of the coefficients in equation (2) is different from zero. The statistic is based on the distribution:

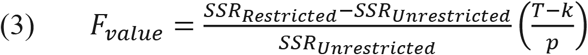

Where, *SSR_Restricted_* is the sum of square residuals of the model which take into account only self-previous observation of the random variable (Equation (1)), and *SSR_Unrestricted_* is the sum of square residuals of the other model which also takes into account the previous observation of the second random variable (Equation (2)). *T* is the sample size (number of observations in the time series used for prediction), *p* is the number of variables which was removed from the unrestricted model, in our case, the lag order, and *k* is the number of variables, in our case, twice the lag order. The null hypothesis is rejected when the *F_value_* is larger than the *F* statistic (i.e., F’s critical value) to conclude that *GC_x_j_ −>_x_i___*. We derived the p-value from the F-statistic instead of directly using the F statistic to set up a global acceptance threshold.

### Calculating the transmission and receiver scores

The transmission and the receiver scores were calculated as the probability for an outgoing (respectively, ingoing) edge at topological distance ≤ 2 (nearest and next-to-nearest neighbor), where the topological distance was calculated using the Delaunay triangulation. These neighborhood sizes were determined for sufficient observations for statistics, and the expected short-range communication between the cells. The transmission (denoted Tr) and receiver (denoted Re) scores were calculated independently for each cell *c_i_* using its cell neighbours at topological distance one and two 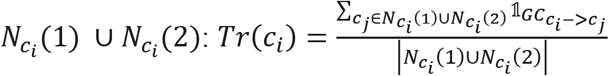 and 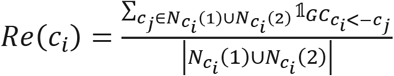.

Importantly, we treat each cell as an independent observation thus characterizing the role of each cell in the multicellular network without specifically committing on the exact network edges. To fix spurious edges due to multiple hypothesis testing we applied the strict Bonferroni correction that defines the edge significance threshold based on the number of edges considered ^63^. In our case, with a significance threshold of 0.05 and n - number of potential edges we get a new significance threshold of 0.05/n. Edges passing these strict statistical tests are termed *GC edges*.

### Partitioning the normalized transmission-receiver space

The transmission and receiver scores of each cell were normalized across the population to allow direct comparison of single cell heterogeneity. We calculated the receiver and transmission z-score for each cell *c_i_*, the variation from the mean in units of standard deviations: Tr_norm(ci) = (Tr(ci)-μ)/σ, where μ is the mean transmission score across the population, and σ is the standard deviation. The same normalization was applied for the receiver score. Kernel Density Estimation ^64^ was used for the visualization of the 2-dimensional normalized transmission and receiver score space (Fig. 2D, Fig. 3E). We partitioned the normalized transmission-receiver space to five regions, and assigned each cell to one of these regions. Individual: transmission and receiver z-score < −0.5. Common: transmission z-score in the range of −0.5-0.5 and receiver z-score < 0.5 or receiver z-score in the range of −0.5-0.5 and transmission z-score < 0.5. Leader: transmission z-score > 0.5 and receiver z-score < 0.5. Follower: receiver z-score > 0.5 and transmission z-score < 0.5. Leader: transmission and receiver z-score > 0.5. The z-score threshold of 0.5 was selected to maintain sufficient number of cells in each role for statistical analysis.

### Measuring heterogeneity

We defined two measures for heterogeneity and applied it to each biological replica of the “step” experiments. First, the ratio between the number of hub cells and the number of individual cells (termed *hub/individual ratio*). Second, the bulk homogeneity, which was calculated as the Earth Mover’s Distance (EMD) ^31–33^ between the experimentally observed and the uniform distribution for all cells that hold |transmission score + receiver score| < 1 (~40% of the cells), thus considering the combined GC outdegree and indegree scores for each cell. The bulk homogeneity was calculated based on observed and uniform distributions that were binned to 20 intervals to cover the [-1,1] range. Importantly, hub/individual ratio and bulk homogeneity were independent and complementary measures for heterogeneity because the cells considered for the bulk homogeneity calculation did not include hubs or individuals.

To verify that these heterogeneity measures are local, i.e., dependent on the spatial organization of the cells, we performed permutation tests. For hub/individual ratio we randomly switched the location of each hub and individual cell with another cell in topological distance > 2. This was followed by re-calculating the transmission and receiver scores for all the cells that were involved in this switching defining a new hub/individual ratio value. This process was repeated multiple times (Fig. S8A). For bulk homogeneity, we shuffled the cells’ spatial locations and recalculated the bulk homogeneity based on the new spatial ordering. This was followed by correlating the bulk homogeneity with the synchronization measures, which were independent of the cells’ spatial ordering, across all “step” experiments. This process was repeated ten times and compared to the series of bulk homogeneity scores of the step experiments with the original location of the cells (Fig. S8D-E).

### Measuring information flow

*GC edge probability* was defined as the probability of a GC edge in the experiment. This was calculated as the ratio between the total number of GC edges and the total number of potential edges in the experiment. Because GC edge probability is a proxy for the information flow within the multicellular network, we also used the term *information flow* to refer to the GC edge probability in the manuscript text.

### Earth Mover Distance (Wasserstein distance) between the cycles

The Earth Mover’s Distance (EMD) ^31–33^ calculates dissimilarities between distributions as the minimum ‘energy’ required to shift one distribution to the other distribution. We used EMD to calculate the rate of change in the information flow throughout the cycles. To compare two cycles we calculated the EMD between the corresponding two-dimensional transmission-receiver scores cell distributions. We discretized the distribution to 20×20 bins in the range of [−4, 4] in the transmission and receiver axes.

### Enrichment factor of cellular state transitions

We calculated the enrichment factor, the fold change in the observed transition probabilities of single cells from one state (functional role) to another cell state in consecutive cycles in relation to a null model derived from the expected transitions based on the marginal distribution of cells’ functional roles. We first constructed the accumulated transition matrix, where the (i,j) bin holds the total number of transitions of single cells between state i and state j in consecutive cycles, *c*, *c+1* observed throughout an experiment.

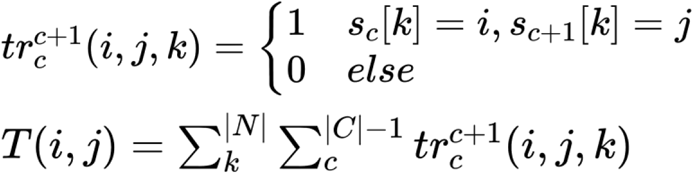

Next, we divided each row *r* of the accumulated matrix by the sum of the row values of *r* to compute the Markov transition matrix (Fig. S14 top-left):

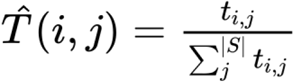

To calculate the enrichment factor, the fold changes in switching from one state to another compared to the expected probability from a null model assuming random transitions drawn from the marginal state distribution, we calculated the expected transition matrix *e_i,j_* (Fig. S14 bottom-left) and divided the Markov transition matrix bins the corresponding bins in the expected matrix (Fig. S14 right).

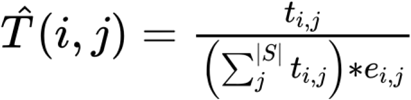

### Measuring cell memory

To measure the cell memory we calculated the Pearson correlation of the cells transmission or receiver scores between consecutive cycles with step Δt (Δt = 1 in Fig. 4A, Δt ≥ 1 in Fig. S12B). We evaluated the significance of our results using a permutation test by shuffling the cells’ spatial locations with over 1000 permutations. The permutation test was performed by concatenating the vector scores of the cycles c, c+Δt, shuffling the values, splitting back to two vectors, and calculating the absolute Pearson correlation. The p-value is set as the fraction of permutations where the shuffled correlation surpassed the observed experimental correlation.

### Activation Time

The activation time of a cell in a given cycle is the time where its calcium dynamics exceeds a threshold value of *γ* within a cycle. The threshold is parametrized by *δ* in the range of 0.1, 0.2 or 0.3 from the calcium dynamics range - the initial value subtracted from the maximal value within the cycle.

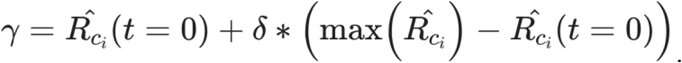

The initial time was shifted by 60 seconds (30 frames) from the onset of the cycle to the time where the mean value of the cells’ calcium dynamics is zero to ensure that the single cell calcium signal is on the rise for the vast majority of the cells.

### Correlating the topological distance between pairs of cells to their GC-edge probability

In Fig. 5B we correlated the topological distance to the corresponding GC edge probabilities. For each topological distance, for each cell, we randomly selected ten (or less in topological distances with smaller numbers) cells and calculated the GC statistical test for each cell pair in both directions. We evaluated the critical value (i.e., p-value correction) using FDR, to correct for multiple hypothesis testing. Finally, we calculated the probability for GC significant edge as the total number of significant edges divided by total GC tests performed.

### Data

N = 47 biological replicates for the “step” experiments: n = 6 (0.1 Pa), n = 13 (0.2 Pa), n = 8 (0.6 Pa), n = 10 (1 Pa), n = 10 (1.6 Pa). N = 6 biological replicates of the “cycle” experiments: n = 2 (0.1 Pa) and n = 4 (2 Pa).

### Software and data availability

The source code and sample data are publicly available in the following link: https://github.com/zamiramos/cyclic-calicum-synchronization

## Funding and Acknowledgements

This work was supported by the Israeli Council for Higher Education (CHE) via Data Science Research Center, Ben-Gurion University of the Negev, Israel (to AZ), and National Institute of General Medical Sciences grant R35GM138179 (to BS). GL is supported by National Science Foundation Grant PHY-1844627. Part of this research was conducted at the Northwest Nanotechnology Infrastructure, a National Nanotechnology Coordinated Infrastructure site at Oregon State University which is supported in part by the National Science Foundation (grant NNCI-1542101) and Oregon State University. We thank Kevin Dean for critically reading the manuscript.

## Author Contribution

AZ and BS conceived the study. GL and KC designed the experimental assay and performed all experiments. Amos Z developed analytic tools, analyzed, and interpreted the data. AZ, BS and Amos Z drafted the manuscript. BS and AZ mentored the authors. All authors wrote and edited the manuscript and approved its content.

## Supplementary Figures

**Figure S1:**
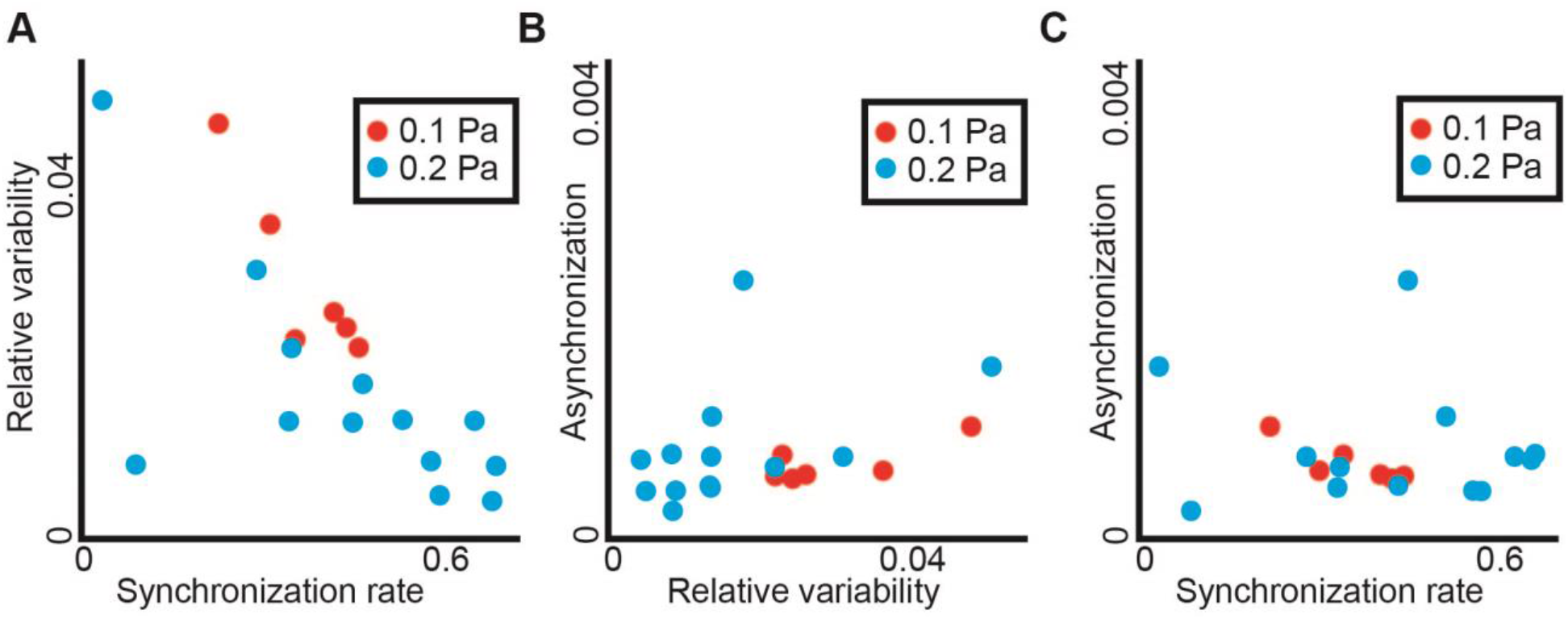
Correlation of synchronization measures. (**A**) Synchronization rate versus relative variability. Pearson correlation = −0.69, p-value = 0.0011. (**B**) Relative variability versus asynchronization. Pearson correlation = 0.345 (not significant). (**C**) Synchronization rate versus asynchronization. Pearson correlation = −0.15 (not significant). For all panels, each observation represents a biological replica. N = 19 replicates: n = 6 experiments with pressure of 0.1 Pa, n = 13 with 0.2 Pa.

**Figure S2:**
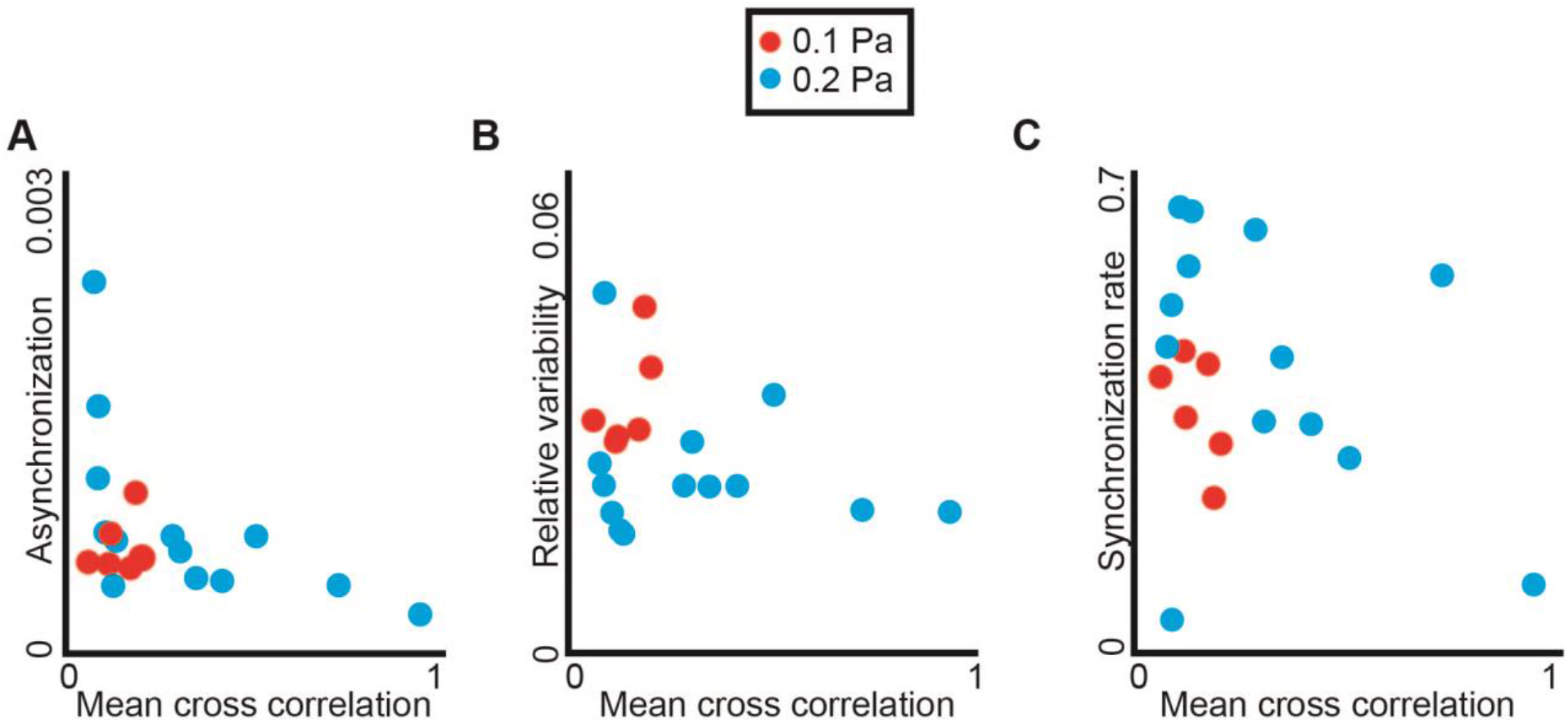
Correlation between the mean neighboring cell-cell temporal correlation in calcium dynamics and multicellular synchronization measures. (**A**) Asynchronization rate versus mean cross correlation. Pearson correlation = −0.46, p-value = 0.046. (**B**) Relative variability versus mean cross correlation. Pearson correlation = −0.267 (not significant). (**C**) Synchronization rate versus mean cross correlation. Pearson correlation = −0.287 (not significant). For all panels, each observation represents a biological replica. N = 19 replicates: n = 6 experiments with pressure of 0.1 Pa, n = 13 with 0.2 Pa.

**Figure S3:**
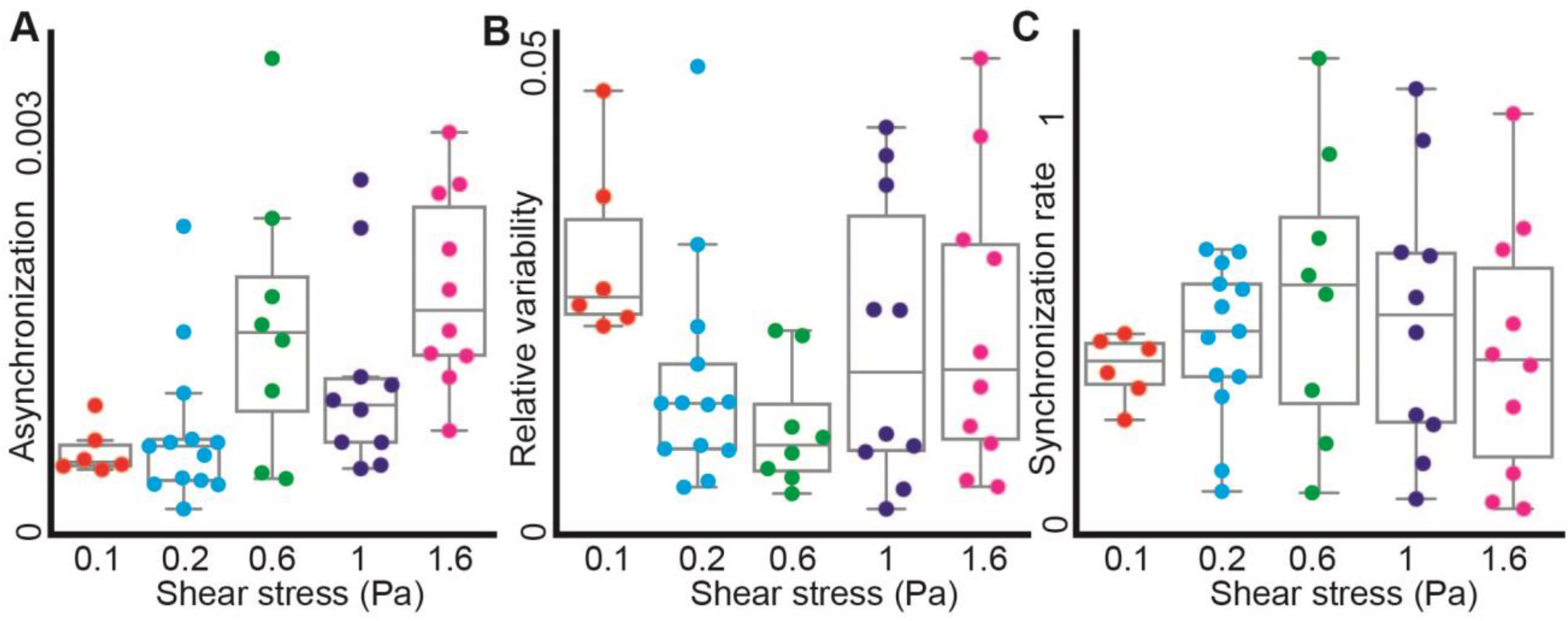
Multicellular synchronization for increasing shear stress levels. (**A**) Experiments at low shear stresses of 0.1-0.2 Pa are more synchronized compared to higher shear stress levels of 0.6-1.6 Pa. P-value < 0.0001 using the non-parametertic Wilcoxon ranksum test. (**B**) Relative variability. No clear trend. (**C**) Synchronization rate, no clear trend. For all panels, each observation represents a biological replica. N = 47 biological replicates: n = 6 (0.1 Pa), n = 13 (0.2 Pa), n = 8 (0.6 Pa), n = 10 (1 Pa), n = 10 (1.6 Pa).

**Figure S4:**
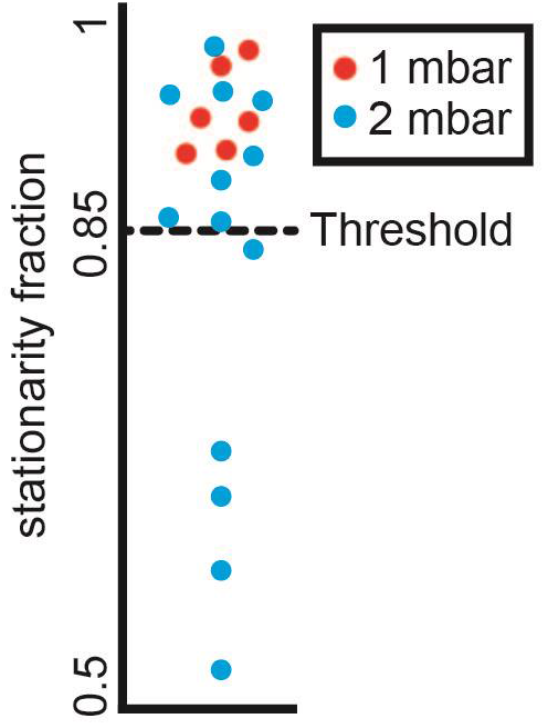
Fraction of cells passing stationarity test per experiment. Each observation represents a biological replica. N = 19 biological replicates: n = 6 (0.1 Pa), n = 13 (0.2 Pa). In 14/19 experiments at least 85% of the cells passed both the KPSS and the ADF stationary tests with significance of below 0.05 (see Methods).

**Figure S5:**
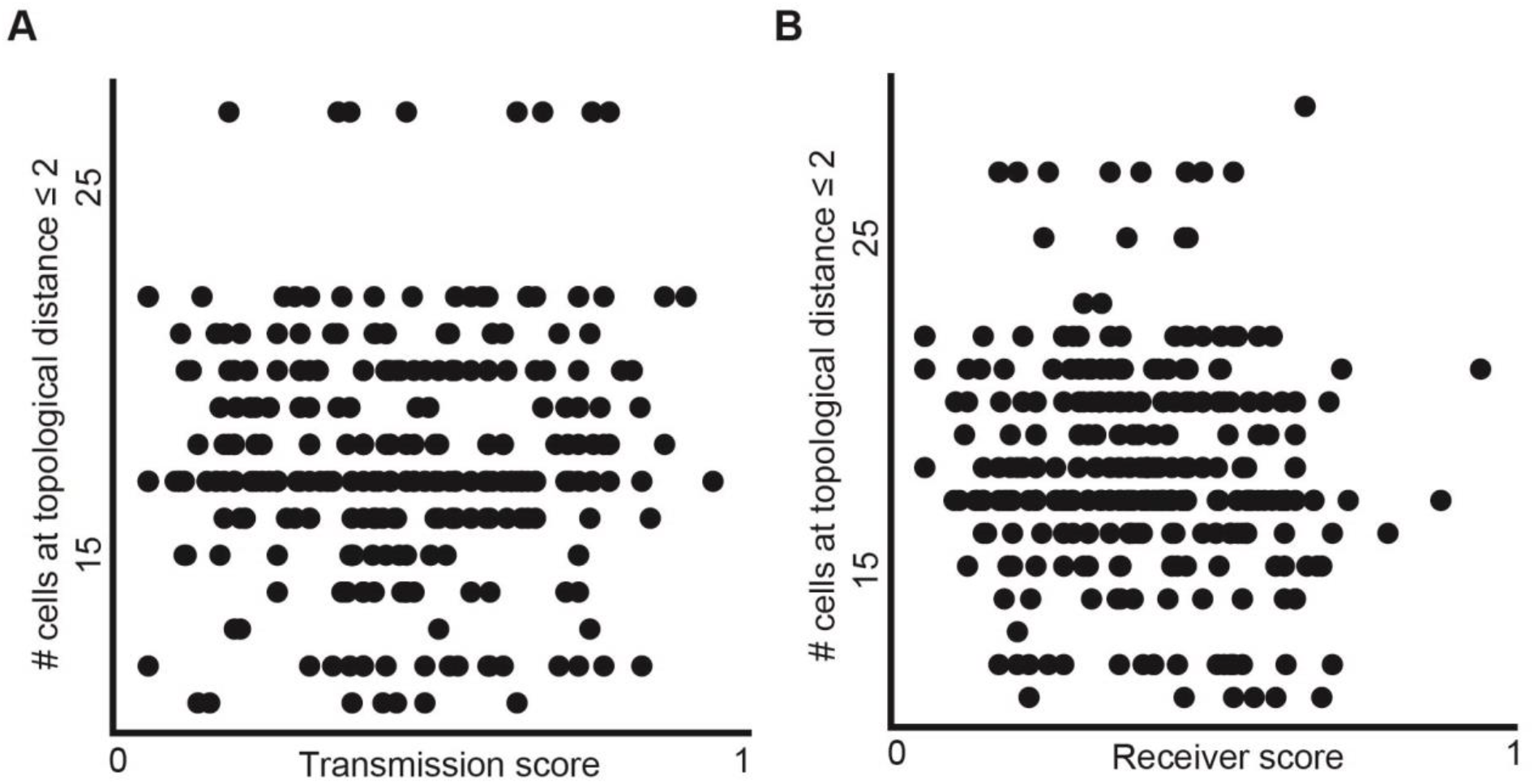
The number of neighbors at topological distance ≤ 2 is not correlated with the transmission or receiver scores. N = 295 cells from one experiment. (**A**) Transmission score. Pearson coefficient = 0.04, p-value = 0.47. (**B**) Receiver score. Pearson coefficient = −0.09, p-value = 0.11.

**Figure S6:**
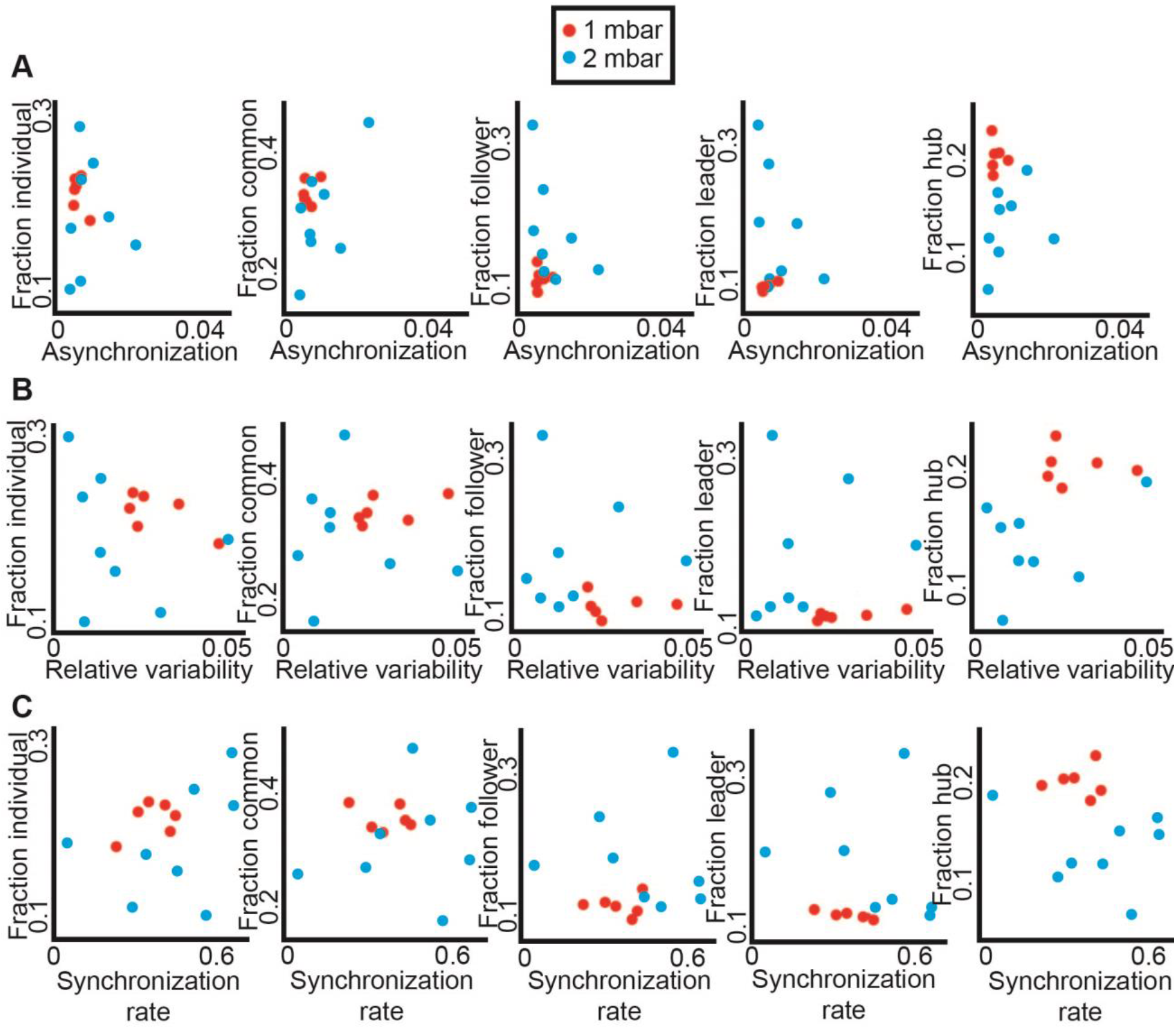
Fraction of cells in each role is not associated with any synchronization measure. (**A**) Asynchronization was not correlated to the fraction of any specific role. Left to right: Individual: Pearson coefficient = −0.13, p-value = 0.66; Common: Pearson coefficient = 0.48, p-value = 0.08; Follower: Pearson coefficient = −0.20, p-value = 0.48; Leader: Pearson coefficient = −0.15, p-value = 0.61; Communication hub (“hub”): Pearson coefficient = −0.09, p-value = 0.75. (**B**) Relative variability was not correlated to the fraction of any specific role. Left to right: Individual: Pearson coefficient = −0.196, p-value = 0.5; Common: Pearson coefficient = 0.045, p-value = 0.88; Follower: Pearson coefficient = −0.184, p-value = 0.53; Leader: Pearson coefficient = −0.053, p-value = 0.86; Communication hub (“hub”): Pearson coefficient = 0.476, p-value = 0.08. (**C**) Synchronization rate was not correlated to the fraction of any specific role. Left to right: Individual: Pearson coefficient = 0.32, p-value = 0.27; Common: Pearson coefficient = 0.06, p-value = 0.84; Follower: Pearson coefficient = 0.04, p-value = 0.90; Leader: Pearson coefficient = −0.12, p-value = 0.67; Communication hub (“hub”): Pearson coefficient = −0.31, p-value = 0.29. For all panels, each observation represents a biological replica. N = 14 replicates: n = 6 (0.1 Pa), n = 8 (0.2 Pa).

**Figure S7:**
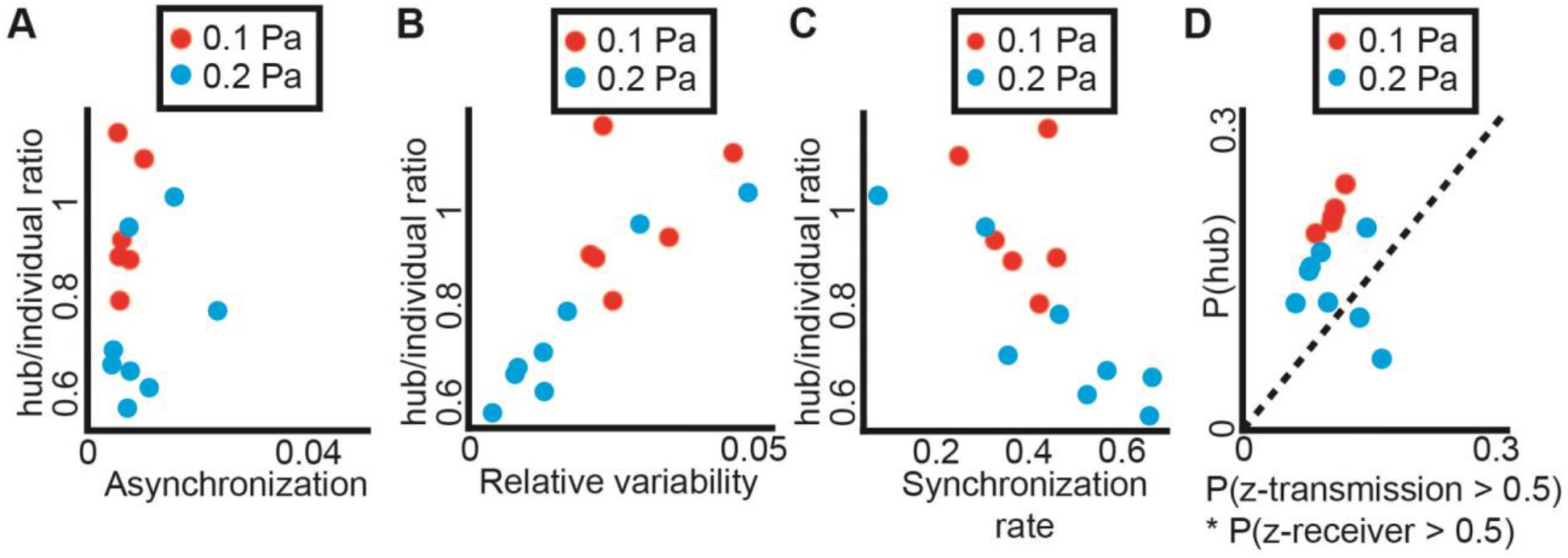
Correlation between synchronization measures and the hub/individual ratio, defined as the ratio between the fractions of cells in the “hub” versus “individual” roles. (**A**) Asynchronization and the hub/individual ratio were not correlated. Pearson correlation = 0.05, p-value = 0.86. (**B**) The hub/individual ratio was correlated with the relative variability. Pearson correlation = 0.8227, p-value = 0.0003. (**C**) The hub/individual ratio was negatively correlated to the synchronization rate (lower rate implies faster synchronization). Pearson correlation = 0.7203, p-value = 0.0037. (**D**) Communication hubs were enriched beyond their expected probability. p-value = 0.0076 via Wilcoxon rank-sign test on P(hub)-P(z-transmission)*P(z-receiver). For all panels, each observation represents a biological replica. N = 14 replicates: n = 6 (0.1 Pa), n = 8 (0.2 Pa).

**Figure S8:**
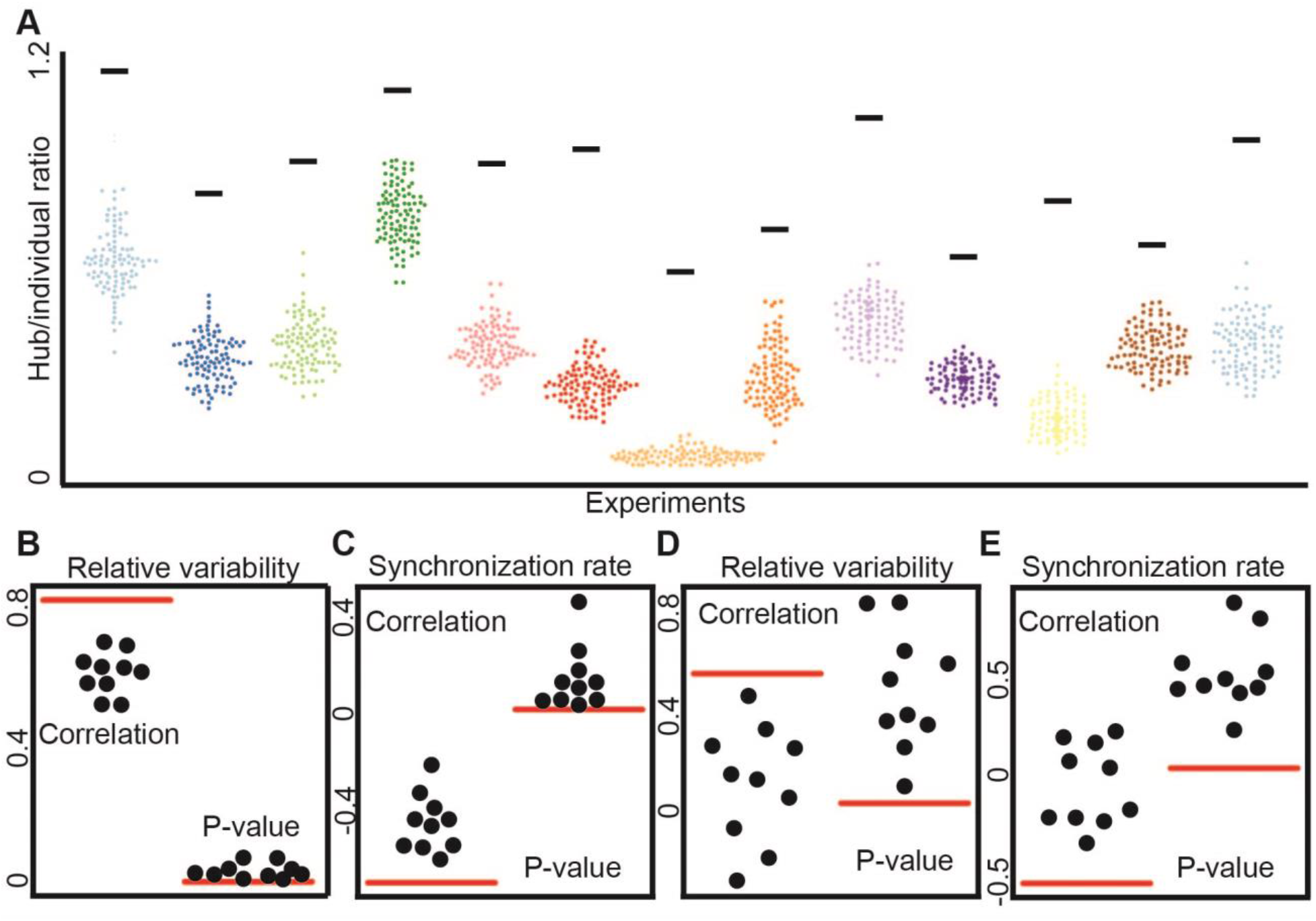
Heterogeneity measures depend on the cells’ spatial organization and are thus local measures. Horizontal lines represent experimental observations, data points represent permutation test results (Methods). **(A)** Hub/individual ratio depends on the cells’ spatial organization. Each color represents a different experiment. Each dot represents one permutation test simulation. N =100 permutation test simulations were performed for each experiment. **(B-E)** Correlations between heterogeneity and synchronization measures. In each panel, the left side shows the Pearson correlations across N = 14 biological replicas, and the right side shows the corresponding p-values. Ten dots per plot represent different simulated observations that are compared to the experimental observation (red horizontal line). Each dot represents the correlation/p-value between the synchronization and the corresponding simulated heterogeneity across experiments. The synchronization measures are global properties that do not change in the simulated permutations. We did not assess the asynchronization measure because it did not correlate with any of the heterogeneity measures. **(B-C)** Correlation between the hub/individual ratio and synchronization measures. (**B**) Correlation to relative variability (observed Pearson correlation = 0.8227, p-value = 0.0003). **(C)** Correlation to synchronization rate (observed Pearson correlation = −0.7203, p-value = 0.0037). **(D-E)** Correlation between bulk homogeneity and synchronization measures. **(D)** Correlation to synchronization rate (observed Pearson correlation = 0.5582, p-value = 0.0380). **(E)** Correlation to synchronization rate (observed Pearson correlation = −0.5643, p-value = 0.0355).

**Figure S9:**
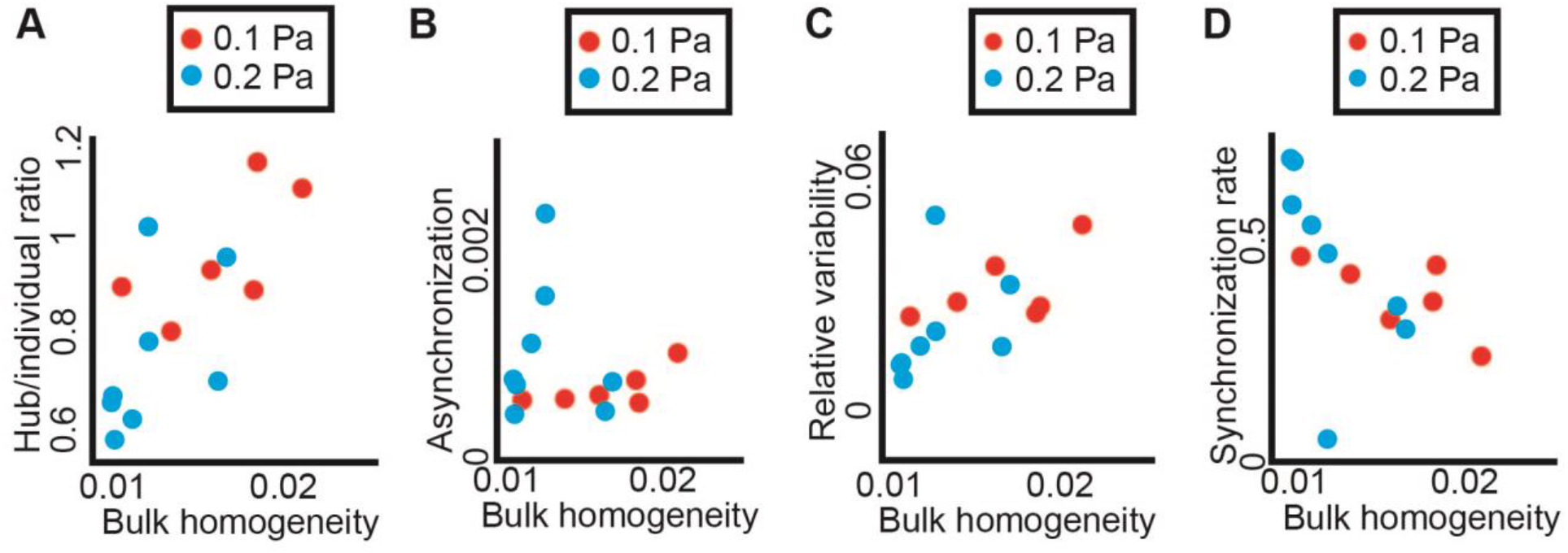
Bulk homogeneity association with other heterogeneity and synchronization measures. (**A**) Bulk homogeneity is correlated to the hub/individual ratio. Pearson correlation = 0.72, p-value < 0.004. (**B**) Bulk homogeneity is not correlated to the asynchronization. Pearson correlation = 0.13, p-value = 0.64. (**C**) Bulk homogeneity is negatively correlated with the relative synchronization. Pearson correlation = 0.56, p-value = 0.038. (**D**) Bulk homogeneity is negatively correlated with the synchronization rate. Pearson correlation = −0.56, p-value < 0.036. For all panels, each observation represents a biological replica. N = 14 replicates: n = 6 (0.1 Pa), n = 8 (0.2 Pa).

**Figure S10:**
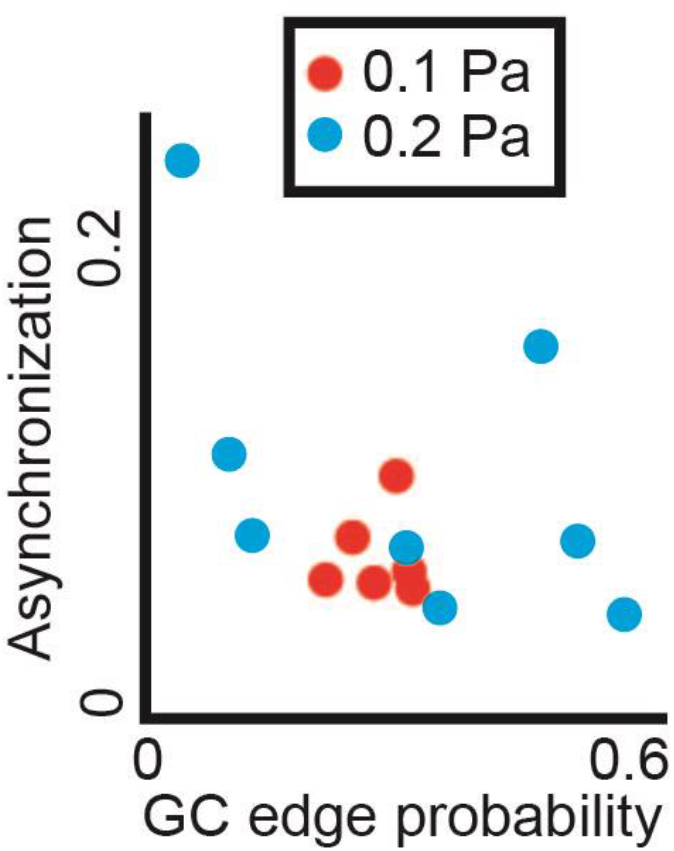
Asynchronization was not associated with the Granger Causality edge probability - the fraction of significant GC tests in an experiment. N = 14, Pearson correlation = −0.42, p-value = 0.14.

**Figure S11:**
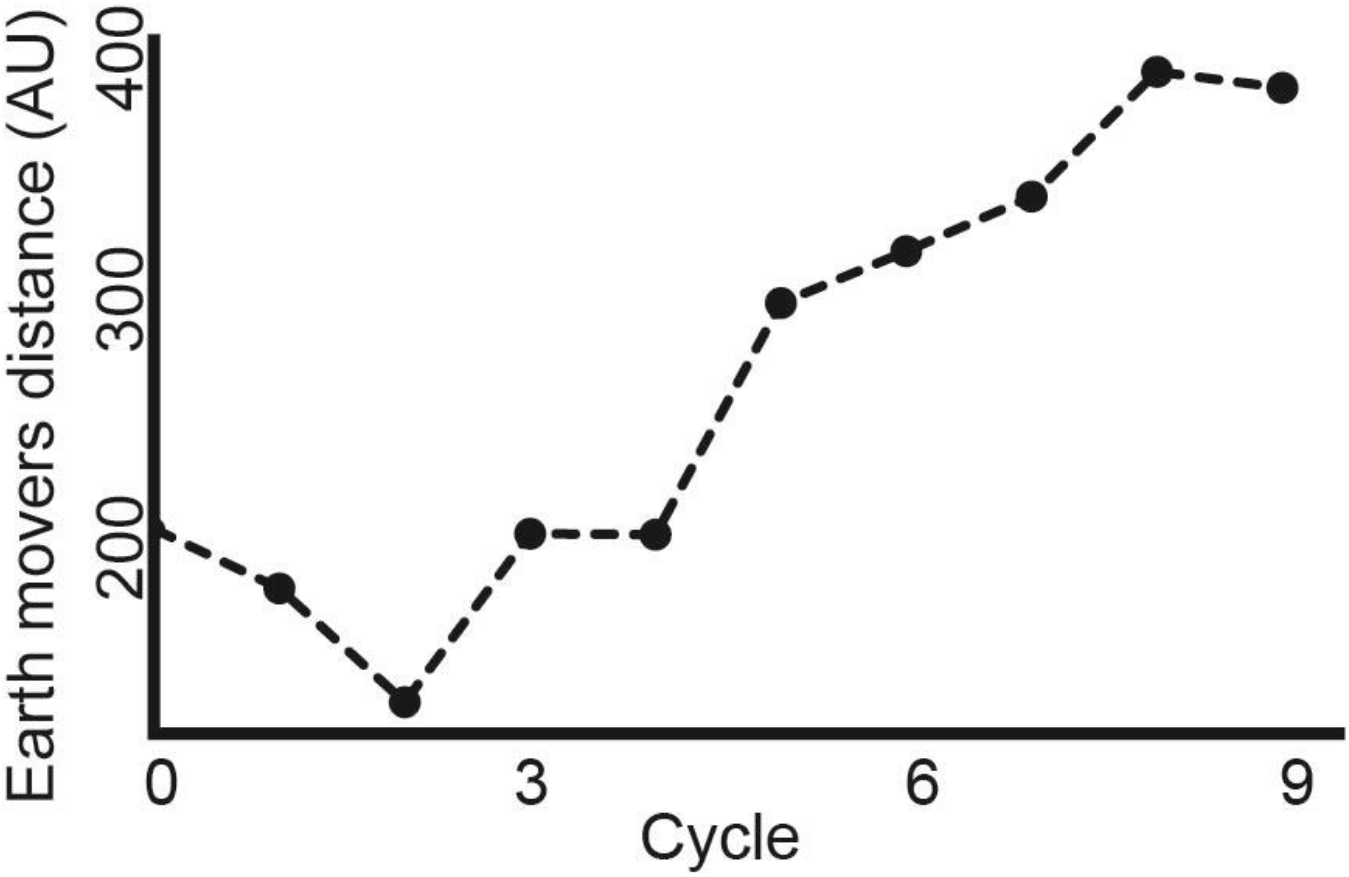
The cells’ roles distribution change rate is increased over the cycles. Each dot represents the Earth mover’s distance (EMD) between the cell’s roles distribution of two cycles (*i* and *i+3; for i = 0 .. 9*) using the Wasserstein metric. (Methods).

**Figure S12:**
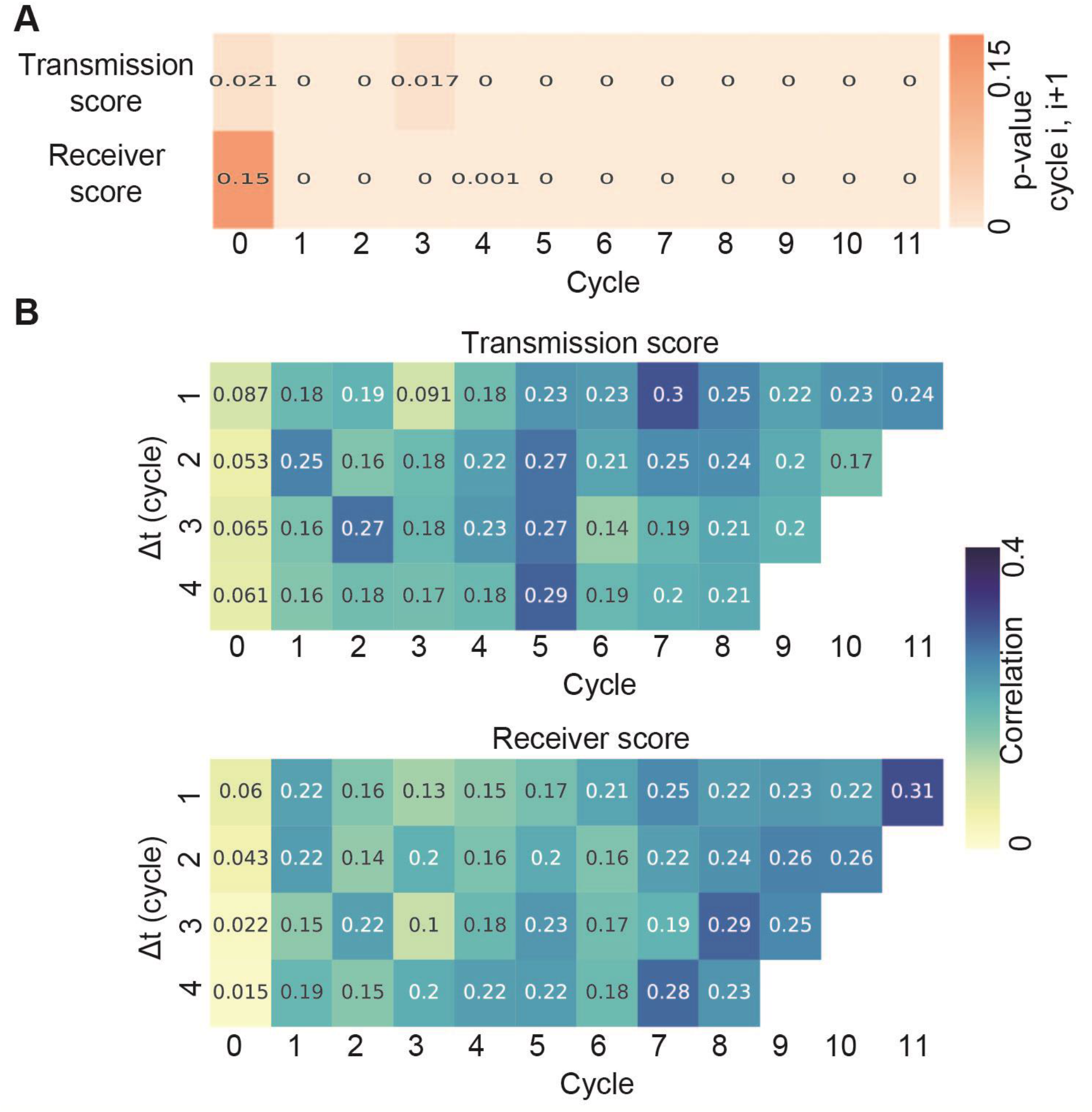
Single cell communication memory. (**A**) P-value of the correlation between the cells’ transmission/receiver scores over consecutive cycles, values correspond to the correlations in Fig 4A. (**B**) Longer term memory. The temporal correlations between single cells’ transmission (top) or receiver (bottom) scores with time gaps of 1-4 cycles (y-axis).

**Fig. S13:**
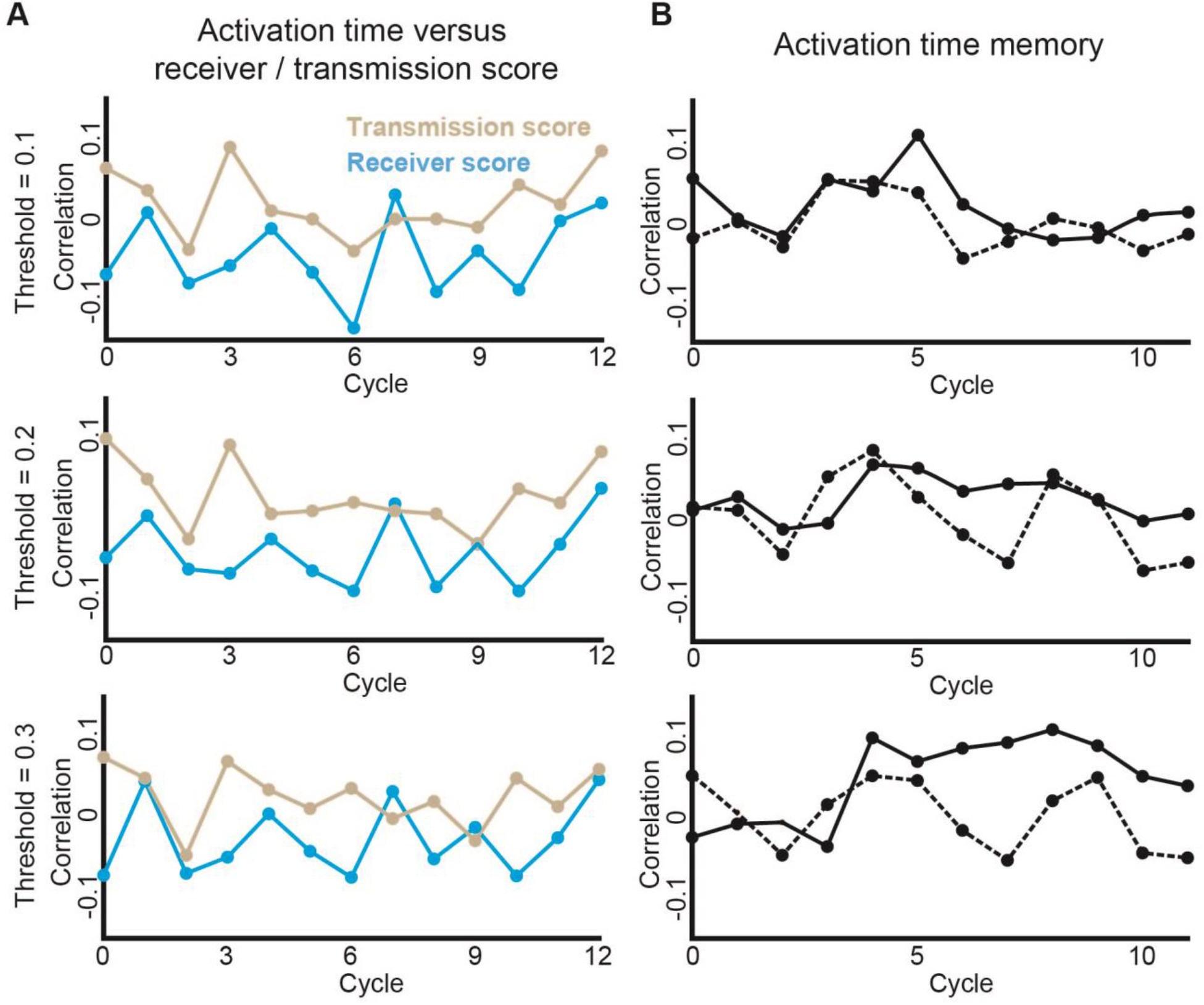
Single cell activation time. A cell’s activation time was defined as the time where the cell’s calcium dynamics exceed *δ* = 0.1-0.3 (top-to-bottom) from it’s maximal signal, see Methods for full details. **(A)** Cells’ activation time was not correlated with the transmission or the receiver score. **(B)** Cells’ activation time was not correlated across consecutive cycles (solid lines). The dashed lines were the calculated correlation after shuffling the cells in the next cycle.

**Figure S14:**
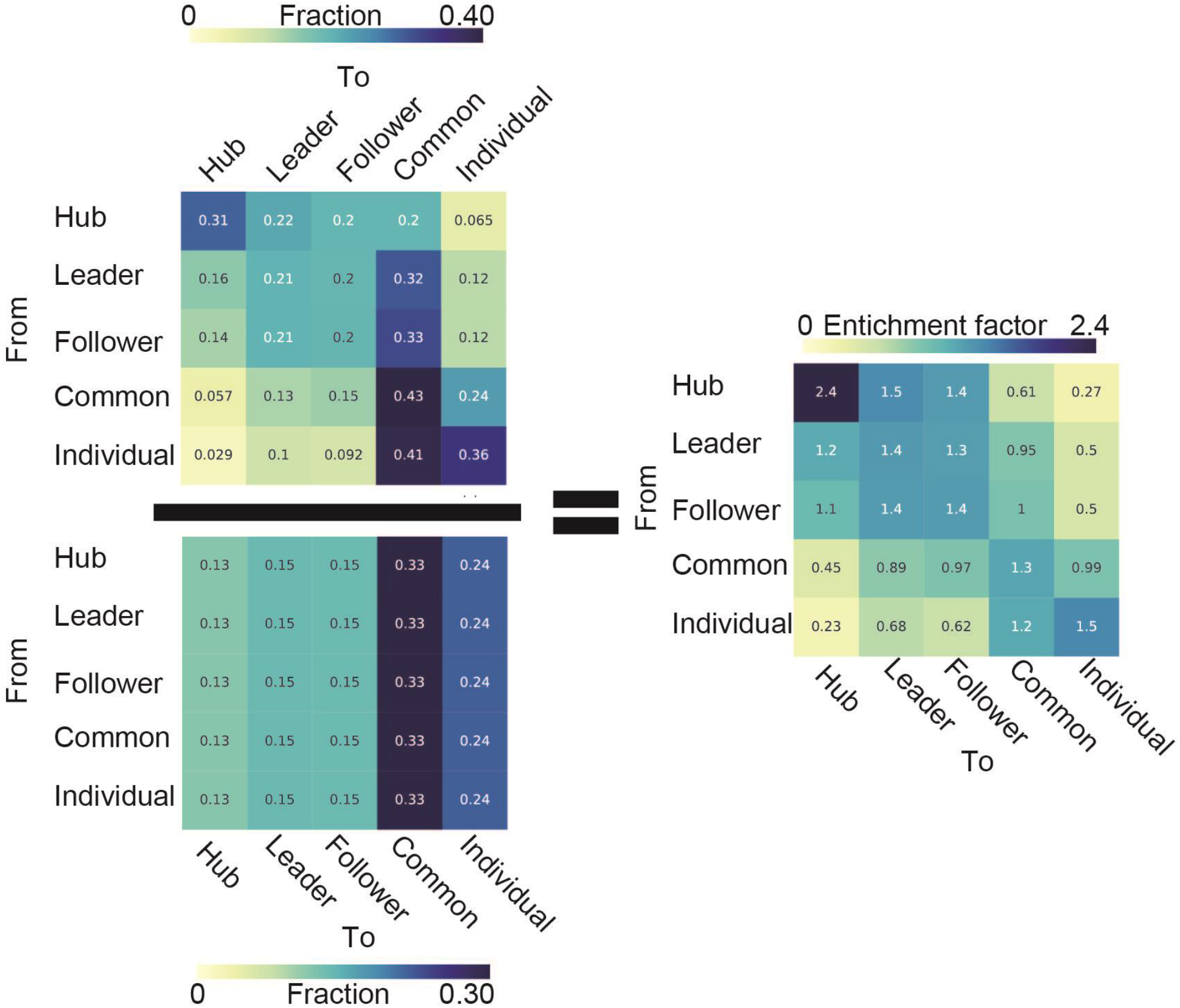
Calculating the enrichment factor of cellular state transitions. Top left: Markov transition matrix based on the observed single cell role transition over the cycles. Bottom left: The expected transition probability between two roles based on the cells’ roles frequency over the cycles. Right: The enrichment transition matrix was followed by normalizing (scalar division) the Markov transition matrix with the expected transition matrix. The enrichment factor from hub to hub is higher than any other enrichment factor. (see only transitions above the expected, Fig. 4D).

## Supplementary Video Legends

**Video S1**: A video of a representative “step” experiment with shear stress of 0.2 Pa.

**Video S2**: A video of a cyclic experiment of shear stress of 0.2 Pa.

**Video S3**: A video depicting the transmission score of a representative “cycles” experiment with shear stress of 0.2 Pa. Each region represents a single cell.

**Video S4**: A video depicting the receiver score of a representative “cycles” experiment with shear stress of 0.2 Pa. Each region represents a single cell.

**Video S5**: A video depicting the evolution of the transmission and receiver scores of a representative “cycles” experiment with shear stress of 0.2 Pa. Showing is the kernel density estimate plot visualization of the normalized transmission and receiver score over the cycles. Partitioning of the z-score normalized transmission-receiver space to five regions (blue dashed lines), each cell (yellow dot) was assigned to a state or role (red text) according to the region they resided at.

**Video S6**: A video depicting the single cell evolution in communication properties of a representative “cycles” experiment. Each region represents a single cell and the communication roles are color coded.

